# Stepwise evolution of a butterfly supergene via duplication and inversion

**DOI:** 10.1101/2021.12.06.471392

**Authors:** Kang-Wook Kim, Rishi De-Kayne, Ian J. Gordon, Kennedy Saitoti Omufwoko, Dino J. Martins, Richard ffrench-Constant, Simon H. Martin

**Author notes:** these authors contributed equally (Kang-Wook Kim, Rishi De-Kayne).

## Abstract

Supergenes maintain adaptive clusters of alleles in the face of genetic mixing. Although usually attributed to inversions, supergenes can be complex, and reconstructing the precise processes that led to recombination suppression and their timing is challenging. We investigated the origin of the BC supergene, which controls variation in warning colouration in the African Monarch butterfly, *Danaus chrysippus*. By generating chromosome-scale assemblies for all three alleles, we identified multiple structural differences. Most strikingly, we find that a region of >1 million bp underwent several segmental duplications at least 7.5 million years ago. The resulting duplicated fragments appear to have triggered four inversions in surrounding parts of the chromosome, resulting in stepwise growth of the region of suppressed recombination. Phylogenies for the inversions are incongruent with the species tree, and suggest that structural polymorphisms have persisted for at least 4.1 million years. In addition to the role of duplications in triggering inversions, our results suggest a previously undescribed mechanism of recombination suppression through independent losses of divergent duplicated tracts. Overall, our findings add support for a stepwise model of supergene evolution involving a variety of structural changes.

## INTRODUCTION

Supergenes are clusters of loci at which adaptive combinations of alleles are co-inherited. This can facilitate the maintenance of complex phenotypes under balancing selection[1,2], or link locally adapted alleles in the face of migration[3]. Supergenes are commonly assumed to be associated with a chromosomal inversion polymorphism[4], but detailed reconstruction of some supergenes has revealed more complex architectures, such as multiple adjacent inversions that arose in a stepwise manner[5], similar to the progressive spread of recombination suppression in sex chromosome evolution[6]. Unlike a single inversion, supergenes maintained by more complex mechanisms could protect more than two divergent alleles from recombination. Other forms of genomic rearrangement that disturb local synteny may also contribute to recombination suppression[7,8], but there are few well-characterised cases, and in some cases the precise mechanism of recombination suppression is unknown[9]. One effective approach to explore the structural changes involved in supergene evolution and their relative timing is to compare whole-chromosomal assemblies for each of the supergene alleles[10].

We investigated the evolution of a colour patterning supergene in *Danaus chrysippus*, a dispersive butterfly known as the African Monarch, African Queen or Plain Tiger in different parts of its extensive range. It has bright warning colouration indicating its toxicity[11], and is divided into distinct geographic morphs with partly overlapping ranges[12]. Despite its vast range, genetic differentiation is almost non-existent between distant populations: *F*_ST_ is approximately zero across the genome, with the exception of a few broad peaks, including two that are associated with colour differences[13]. This implies selection for the maintenance of local morphs, possibly driven by learned predator avoidance favouring the most common local morph[14]. Colour variation in the forewing is controlled by the BC supergene on chromosome 15[13,15]. This broad region of suppressed recombination includes two colour patterning loci: B controls brown vs orange variation and localises at the melanin pathway gene *yellow*, and C controls the presence/absence of a black forewing tip and white band. Three divergent BC alleles, which determine three distinct forewing phenotypes, have been described to date[13]. A previous genome assembly of one of the alleles (namely ‘klugii’) revealed that several genes on chromosome 15 had increased in copy number in *D. chrysippus* relative to the outgroup *Danaus plexippus* (the Monarch), suggesting that gene duplications may have been involved in the evolution of the BC supergene[13].

To reconstruct the steps in the evolution of the BC supergene, we generated haplotype-resolved chromosomal assemblies for all three of the divergent alleles using long-read sequencing and trio-binning. We traced the evolution of the various structural changes across the phylogeny to determine the sequence of events. Our findings show that the supergene has evolved through multiple steps over the course of several million years, and was preceded by extensive segmental duplications, which likely triggered recombination suppression.

## RESULTS

### Haploid assemblies of the three BC alleles

We generated four new haploid genome assemblies from two F1 individuals by binning reads according to their parent of origin[16] (see Table S1 for accession numbers and Table S2-S4 for assembly statistics). In all four assemblies, chromosome 15 (chr15) is represented by 1-4 contigs that could be ordered and oriented by eye (Figure S1). To identify the supergene allele represented by each assembly, we used an ‘ancestry painting’ approach based on sequence similarity to representative individuals of known genotype[13]. This confirmed that we had assembled all three alleles, as follows: MB18102MAT = ‘chrysippus’, MB18102PAT = ‘klugii’ and SB211PAT = ‘orientis’ (Figure S2). Unexpectedly, the fourth assembly (SB211MAT) showed mixed similarity to all three alleles, suggesting that it may be a rare recombinant that has not yet been observed in the homozygous state (Figure S2, S3).

Using the same ancestry painting procedure, we also confirmed that a previous assembly (Dchry2.2)[17], represents a ‘pure’ klugii allele (Figure S2), despite having been assembled from diploid reads from a suspected heterozygous individual. We therefore examined the discarded haplotypic contigs from the Dchry2 assembly (‘Dchry2HAP’) and identified the other chr15 haplotype, which represents the chrysippus allele (Figure S2). Therefore, in total, we have generated six assemblies of chr15: two representing the klugii allele, two representing the chrysippus allele, one representing the orientis allele, and one representing a putative recombinant. The two independent assemblies of the klugii and chrysippus alleles allowed us to check for assembly errors. In both cases the independent assemblies of the same chr15 allele were co-linear with no obvious errors detected (Figure S4).

### Multiple structural rearrangements in the evolution of the BC supergene

We predicted that each of the three BC alleles has undergone distinct structural changes resulting in recombination suppression. We therefore first analysed the alignments between each allele and the outgroup *D. plexippus* assembly, assumed to be structurally unaltered from the ancestral state. As predicted, each allele shows multiple structural differences from *D. plexippus* (Figure 1, Figure S1). Most of these differences are unique to one of the three BC alleles, implying that they are derived changes. The structural events involve four adjacent chromosomal regions. The most strikingly rearranged is Region 3 (1.3 megabases [Mb]), which has been duplicated and fragmented multiple times in all three resulting alleles. Many of the duplicated fragments have been inverted and translocated. These multiple duplications result in chr15 being considerably longer in *D. chrysippus* than in *D. plexippus*, with striking variation in length also seen among the BC supergene alleles due to each carrying a different complement of duplicated fragments of Region 3 (Figure 1). Region 3 shows elevated levels of transposable elements (TEs), specifically LINEs, in both the *D. plexippus* and *D. chrysippus* (Figure S5), suggesting that TE activity may have triggered the segmental duplications.

**Figure 1.**
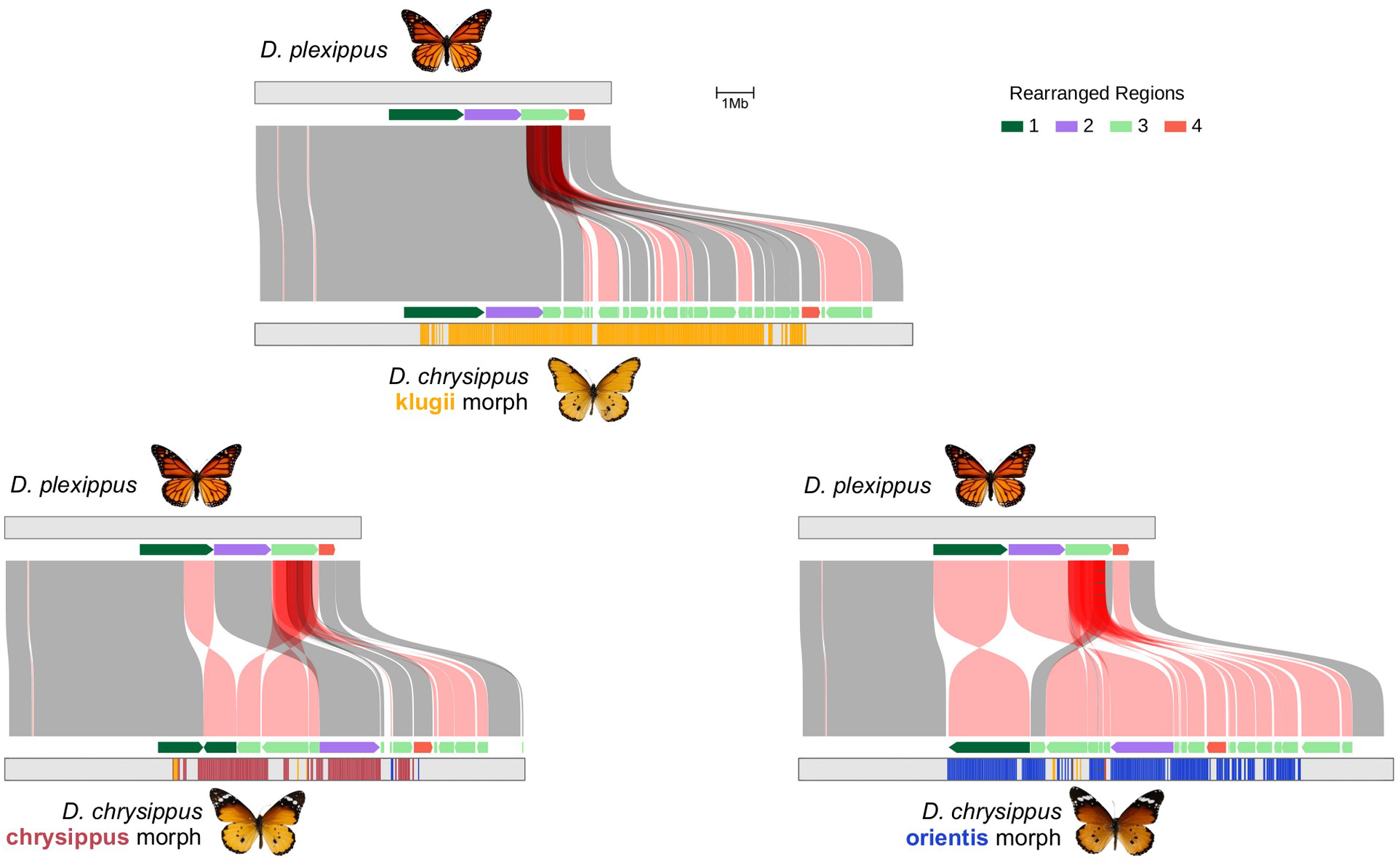
Structural changes in *D. chrysippus* supergene alleles relative to the outgroup. Connecting boxes indicate syntenic blocks (see Methods for details) between the outgroup (*D. plexippus*, MEX_DaPlex assembly) and each of the three BC supergene alleles in *D. chrysippus*. Blocks that are syntenic but in the reverse orientation are coloured red. Partial transparency is used to reveal duplicated syntenic blocks. Coloured arrows indicate the four regions that we identified as having experienced distinct rearrangements in *D. chrysippus*, with the direction of the arrow indicating their relative orientation. The coloured bars indicate ‘ancestry painting’ in 50 kb windows along the *D. chrysippus* chromosomes (see also Figure S2). Regions lacking coloured bars could not be assigned ancestry because they showed high sequence similarity to two or more different morphs.

Although duplications can cause assembly errors if their sequences are too similar, this issue does not appear to have affected our assemblies. Independent assemblies of the same allele show strong similarity (Figure S4), and read depth analyses show no evidence of large collapsed repeats (Figure S6). This suggests that the duplications and repeats are distinct enough to assemble separately, which is also confirmed by the lack of highly similar repeats showing up in whole genome alignments (Figure S4).

The other three Regions - 1 (2 Mb), 2 (1.5 Mb) and 4 (0.4 Mb) - are each inverted in the orientis allele (Figure 1). A portion of Region 1 is also inverted in the chrysippus allele, but appears to share the same right-hand inversion breakpoint. In both the orientis and chrysippus alleles Region 2 is shifted to the right due to upstream insertion of duplicated fragments of Region 3. In all of these simpler rearrangements, either one or both edges are bordered by a duplicated fragment of Region 3. We therefore hypothesised that the duplications of Region 3 may have occurred first and triggered subsequent inversions.

### Multiple duplications deep in the history of the *Danaus* genus

To test the hypothesis that multiple duplications of Region 3 initiated the formation of the supergene, we estimated the copy number of each gene on chr15 in six other *Danaus* species and an outgroup using short read data (see Table S1 for accession numbers). Most of the 476 genes for which we had sufficient data to quantify copy number were present as a single copy in all species. However, a cluster of 10 genes in Region 3 shows high copy numbers in all three *D. chrysippus* morphs as well as three additional species: *Danaus petilia* (Australia), *Danaus gilippus* (Americas), and *Danaus melanippus* (Asia) (Figure S7, Table S5). Our inferred species tree, based on 5954 gene alignments confirms that the multiple duplications of Region 3 originated at the base of subgenus *anosia* (Figure 2B). The tree also resolves previous uncertainty regarding the placement of *D. eresimus*, which groups with *D. plexippus* and *D. erippus*, all of which lack the multiple duplications of Region 3 genes. Four of the duplicated genes are homologous to hemicentin and nephrin (Table S5), members of the immunoglobulin family, but to our knowledge their functions in lepidoptera remain unknown.

**Figure 2.**
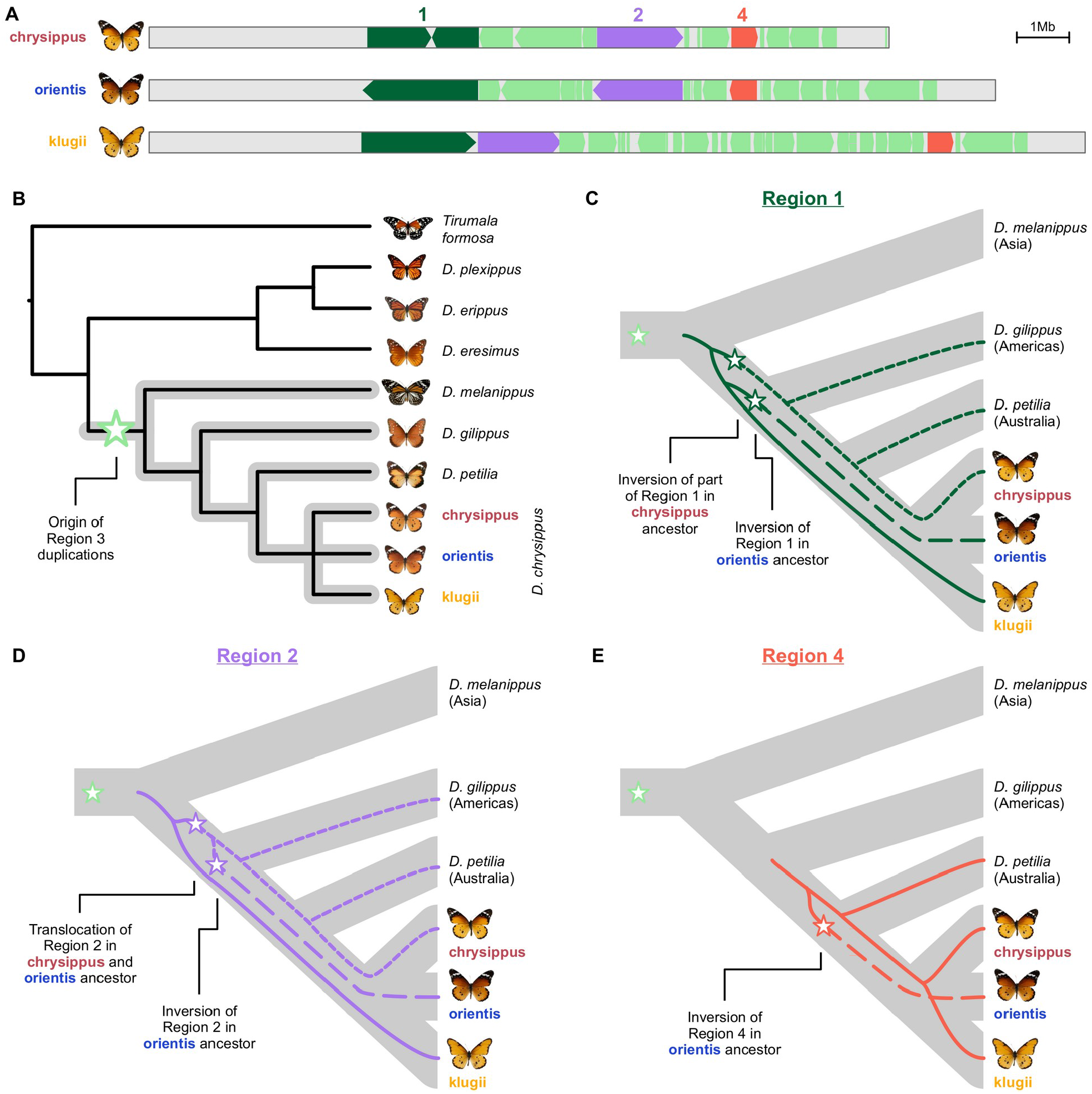
Tracing structural changes across the *Danaus* phylogeny. A. The structure of the three supergene alleles, showing the position and orientation of Regions 1, 2 and 4, which are interspersed by fragments of Region 3. B. Species tree for *Danaus* inferred using ASTRAL, based on 5954 gene alignments. The grey clade indicates all species with elevated gene copy number in Region 3 (Figure S7, Table S5). The green star indicates the inferred origin of the Region 3 duplications. C, D, and E. Inferred genealogies for Regions 1 (including data from 18 genes - 17268 sites for Region 1.2 and 38 - 36576 sites for Region 1.1), 2 (including data from 61 genes - 45214 sites) and 4 (including data from 15 genes - 14127 sites; see Figure S8 for full maximum-likelihood trees). All three genealogies are incongruent with the species branching order. Node ages are therefore positioned to show the most likely origins of the structural variants (indicated by stars) to allow for the observed incongruence. For image attributions see Figure S7.

We estimated a lower bound for the origin of the copy number variation at the base of the *anosia* clade. *d*_a_ (absolute sequence divergence adjusted for estimated divergence in the ancestral species) at putatively-neutral 4-fold degenerate sites between *D. chrysippus* and *D. melanippus* is 8.5% (95% CI = 8.4-8.6). Although we cannot convert this directly to a split time in years as the mutation rates and generation times for this clade are unknown, we can calibrate our estimate according to the estimated split between *Danaus* and *Tirumala* of 12.7 million years ago[18]. *d*_a_ at 4-fold degenerate sites between *D. chrysippus* and *T. formosa* is 14.5% (95% CI = 14.3, 14.6). This gives an estimate of 7.5 million years ago (MYA) as a lower bound for the origin of the Region 3 duplications.

There is consistent variation in estimated copy number for each Region 3 gene among species (Figure 2, Table S5), with the highest copy numbers observed in the *D. chrysippus* morphs. This is most parsimoniously explained by further duplications in the *D. chrysippus* lineage, but there is also evidence for loss of copies (see below). Further work will be required to determine whether all of the duplicated copies are expressed and have phenotypic consequences.

### Structural polymorphisms persist through two speciation events

We next investigated the history of the three simpler rearrangements on chr15 by generating maximum likelihood trees from concatenated coding sequences within each of the three regions involved (38, 18, 61, and 15 genes for Regions 1.1, 1.2, 2, and 4, respectively; Figure S8). The trees for all of the inversions are incongruent with the species branching order (Figure 2C-E), implying that the inversion polymorphisms are ancient and have persisted through speciation events. In Region 1, the first event appears to have been an inversion of part of the region in the ancestor of the chrysippus allele, followed by a separate inversion of the whole of Region 1 in the ancestor of the orientis allele (Figure 2C). Both *D. gilippus* and *D. petilia* are most closely related to the chrysippus allele. This may reflect a scenario in which both *D. gilippus* (found in the Americas) and *D. petilia* (found in Australia) originated through dispersal from North Africa/Asia, where the chrysippus allele is currently found[13]. Using the dating procedure described above, we estimate the split giving rise to *D. gilippus* to 4.1 MYA, implying that the polymorphisms have persisted for at least this long.

Region 2 has also undergone two structural events: an inversion specific to the orientis allele, and a shift (or translocation) caused by insertion of duplicated fragments of Region 3 (Figure 2A). The tree for Region 2 is similar to that for Region 1, indicating recombination suppression among all three supergene alleles pre-dates the emergence of *D. gilippus* and *D. petilia*, who both inherited the chrysippus-like allele (Figure 2D). The node separating the three variants could not be resolved (Figure S8C), but we infer that the translocation shared by both chrysippus and orientis occurred prior to the orientis-specific inversion (Figure 2D). Finally, the tree for Region 4 suggests that the inversion specific to the orientis allele occurred after the speciation event that gave rise to *D. gilippus*, but before the emergence of *D. petilia* (estimated at 1.7 MYA), which inherited the ancestral state (Figure 2E).

### Sequence divergence reveals the landscape of recombination suppression

To understand how the different chr15 rearrangements contribute to recombination suppression in the BC supergene, we examined *d*_*XY*_ (absolute sequence divergence between morphs) and *d*_*a*_ (divergence corrected for within-population diversity) across the chromosome. Elevated sequence divergence is a more reliable indicator of deep coalescence than relative measures such as *F*_ST_[19], and is therefore a useful metric to identify regions of the genome at which recombination has been suppressed between segregating alleles for a long time. For each pair of alleles, clear regions of elevated sequence divergence are detectable, corresponding approximately with the regions that are rearranged between the pair (Figure S9). In Regions 1 and 2, the level of sequence divergence is very high (*d*_*XY*_ ∼ 6%, *d*_*a*_ ∼4%), consistent with these rearrangements having occurred several million years ago, whereas Region 4 shows notably lower divergence, consistent with a more recent inversion (Figure S9). In all three alleles - most notably klugii - some of the duplicated fragments of Region 3 have little or no sequence homology with the other alleles (Figure S9). This implies that some of the paralogs are too divergent to allow cross-mapping of short reads, likely reflecting ancient duplications followed by independent losses of copies in each allele (see Discussion).

## DISCUSSION

By generating chromosome-scale assemblies for all three known alleles of the *D. chrysippus* BC supergene we have uncovered a stepwise process of supergene formation, with a pivotal early role of segmental duplications. Our results suggest two ways in which duplications may contribute to recombination suppression. The first has been described previously: duplicated regions can trigger inversions by promoting ectopic crossovers[20,21]. Meiotic pairing between paralogs can create inverted loops that lead to inversions if crossovers occur on both sides (Figure 3A). In the BC supergene, two of the inverted regions (2 and 4) are bordered on both sides by duplicated fragments, consistent with their involvement in the inversions. Another two inversions involving Region 1 are bordered only on one side by duplicated fragments. The reuse of breakpoints in multiple structural events has been seen elsewhere, and is consistent with a role of duplications or repeats in driving recurrent inversion mutations[10,22–24].

**Figure 3.**
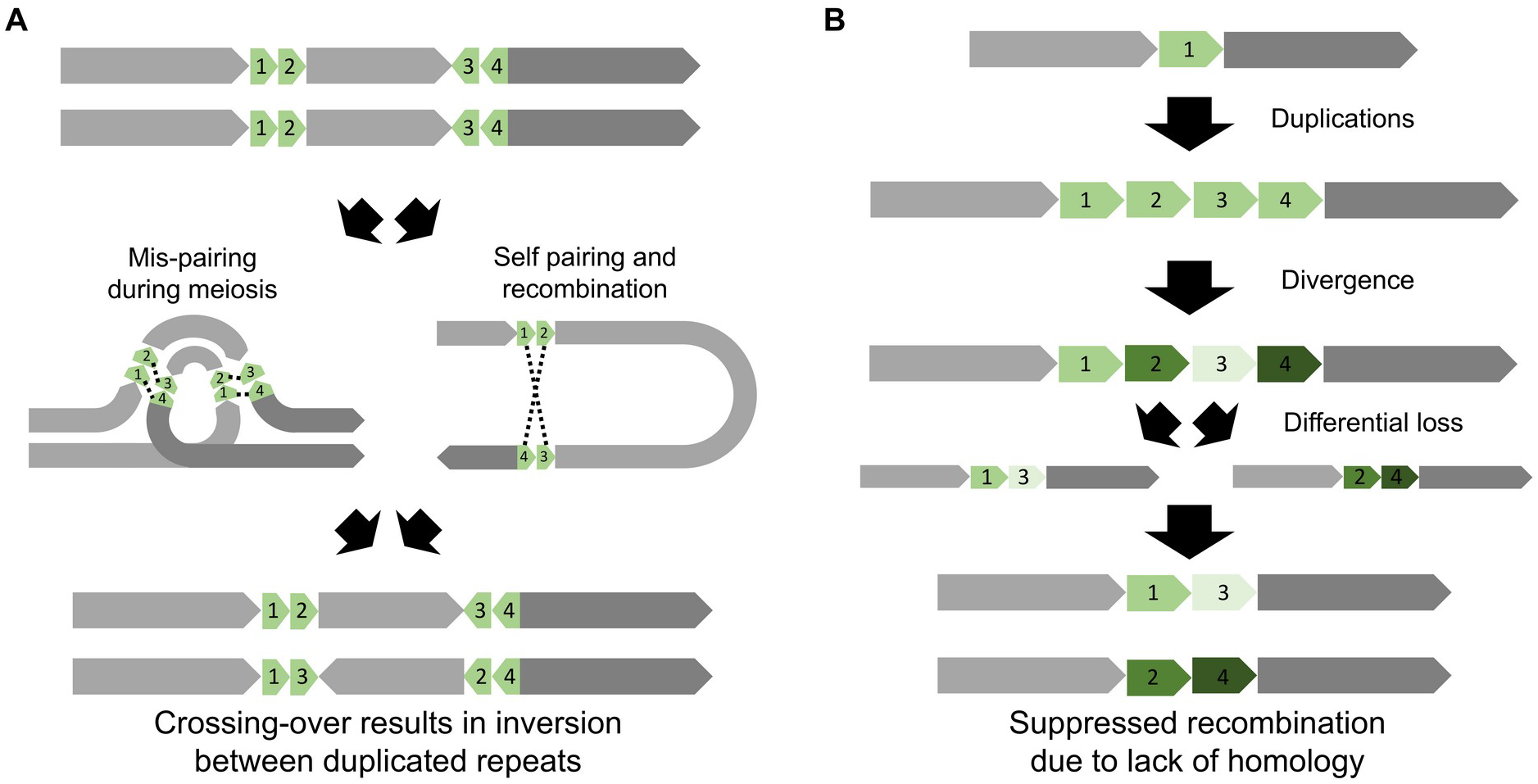
Two mechanisms by which segmental duplications can lead to recombination suppression. A. Ectopic recombination between mis-paired paralogs in different parts of the chromosome can cause an inversion of the intervening region. This mechanism is most relevant when the paralogs are ‘young’ and still share strong sequence similarity. B. Our proposed model of duplication, divergence and differential loss. Over long periods of time, old paralogs may become highly diverged in the sequences. If differential loss of paralogs occurs in different individuals, the remaining copies may be too divergent to pair up and crossover during meiosis.

The second mechanism by which duplications can contribute to recombination suppression has, to our knowledge, not been previously proposed. Our model has three steps (Figure 3B). First, one or more tandem duplications occur and spread to fixation in the population. Second, over a long period, the paralogs become highly diverged through accumulation of mutational differences. Third, independent losses of different paralogs in different individuals leads to the coexistence of haplotypes in a single species that may fail to pair up and form crossovers during meiosis due to lack of homology (Figure S9A). Our model could help to explain the existence of non-recombining regions in the absence of inversions, such as the *doublesex* polymorphism in *Papilio memnon*[9]. It remains to be seen whether duplications and copy number variation are a common feature of supergenes in other taxa.

Following the ancient duplications, four distinct inversions occurred in a stepwise manner to produce the three divergent BC supergene alleles we see today. This stepwise increase in the range of recombination suppression is similar to processes thought to underlie sex-chromosome evolution[6,25]. It is possible that the expansion of the BC supergene has allowed the accumulation of additional co-adapted alleles, and potentially extended its adaptive role to traits beyond colour pattern. The modular nature of the inversions also means that recombination can occur between them without creating unbalanced gametes, and thereby generate recombinant supergene alleles. Indeed, wild individuals frequently show mosaic patterns of ancestry across the BC region[13], and one of the four new assemblies described here appears to be a recombinant (Figure S2). These findings, taken together with recent evidence for extensive gene conversion in inversions in *Drosophila*[26], suggest that the evolution of supergene alleles may be a more dynamic process than previously thought. As long-range sequencing and chromosomal assembly becomes feasible for an increasing number of study systems, we expect that the stepwise mechanisms of recombination suppression described here will prove to be common.

## METHODS

### Samples, processing, and sequencing

To produce haploid assemblies for the three BC alleles, we used a trio binning approach in which long sequencing reads from a diploid sample are binned according to their parent of origin before assembling. We generated two broods (MB181 and SB211) using parents with distinct colour patterns to produce heterozygous F1s carrying two distinct supergene alleles. Specifically, Brood MB181 comprised a father suspected to be homozygous for the klugii allele and a mother suspected to be heterozygous chrysippus/orientis. Brood SB211 comprised a father suspected to be homozygous for the orientis allele and a mother suspected to be heterozygous chrysippus/orientis. From brood MB181, DNA was extracted from a single pupa (MB18102) using the MagAttract HMW DNA Kit from Qiagen (Venlo, Netherlands). PacBio CLR sequencing (Pacific Biosciences, Menlo Park, California, USA) was then carried out using 4 SMRT cells on a PacBio Sequel to produce a total of 34.7 gigabases (Gb) of sequence data. From brood SB211, DNA was extracted from three pupae (SB21101, SB21102, SB21104) and sequenced in a single SMRT cell of a PacBio Sequel II to produce HiFi reads totalling 12.5 Gb of sequence.

In addition to the long-read sequencing of F1s, we performed Illumina sequencing of parents of both broods for trio binning (but used an aunt for MB181 because the mother was lost). We also generated Illumina reads for the F1 pupa of brood MB181 for polishing the assembly, and from an individual of *Tirumala formosa*, to serve as an outgroup. DNA extraction was performed using the Qiagen DNEasy Blood and Tissue KIT. Sequencing was performed using the Illumina NovaSeq 600 platform with 300 cycles and an insert size of 350 bp (see Table S1 for sample information and sequencing coverage).

### Genome assembly and annotation

Although trio binning of long reads requires both parents, we reasoned that a sister of the mother could be used to correctly bin reads from certain chromosomes. Lepidopteran females do not undergo crossing over, so daughters share identical chromosomal haplotypes with maternal aunts for some chromosomes. However, we needed to first confirm that the daughter and aunt shared a first-degree relationship for chr15, the chromosome of interest. To this end, we aligned the Illumina reads to a chromosomal-level *D. plexippus* assembly (‘Dplex_v4’, GCA_009731565,[27]) using BWA v0.7.15 mem[28], converted to BAM format using SAMtools v0.1.18[29], and split by chromosome using BamTools split v2.5.1[30]. We applied IBSrelate[31] to each chromosome separately and confirmed that the F1 and aunt had parent-offspring level relationships for around half the chromosomes, including chr15 (Figure S10).

We used Canu v2.1[32] to perform trio-binning and assembly for MB18102 (Table S2), with estimated genome size of 360Mb and minimum read length of 2 kb. For brood SB211, we had PacBio HiFi sequence data from three offspring (SB21101, SB21102, SB21104). We therefore first binned reads using the *–haplotype* option of Canu, and then pooled the binned reads representing each parent to obtain sufficient read depth for both assemblies. For the assembly, both hifiasm[33] and HiCanu[34] were tested (Table S3). Based on contiguity of chr15-linked contigs (Figure S11), we chose to proceed with the hifiasm assembly for the paternal reads (SB211PAT), and the HiCanu assembly for the maternal reads (SB211MAT).

We assessed assembly completeness using BUSCO v3.1.0[35] with arthropoda_odb9 and insecta_odb9 dataset. As expected, biassed binning occured in the MB181 and more PacBio reads were assigned to father. The paternal assembly (MB18102PAT) therefore shows a high level of duplication whereas the maternal assembly (MB18102MAT) shows a high level of missing BUSCOs (Table S4). However, we assumed that these issues would only affect the chromosomes that do not show parent-offspring relationships with the aunt, and therefore not affect chr15. For all assemblies, redundant (haplotypic) contigs were removed with Purge Haplotigs[36].

Polishing to correct sequencing errors was performed for the two MB18102 assemblies (the SB211 assemblies did not require polishing as they were generated using PacBio HiFi reads). Two rounds of polishing were carried out simultaneously on the MB18102MAT and MB18102PAT assemblies using Illumina data from the assembled F1 individual in the program Hapo-G (allowing the correct mapping of reads to the correct haplotype). Following polishing, each of the MB18102 and SB211 assemblies were repeat masked and annotated using RepeatMasker[37] and BRAKER2[38] pipeline, respectively (as in [17]; using the same Dchry2.2 repeat library and *D. plexippus* protein sets). The transposable element content of the genome was calculated in 50kb windows and the RepeatMasker output using a custom script (Get.TE.Bed.sh; https://github.com/RishiDeKayne/GetTEBed).

### Assembly alignment and rearrangement detection

To identify contigs corresponding to chromosome 15, we performed whole-genome alignment to two chromosomal-level *D. plexippus* assemblies: ‘Dplex_v4’ (GCA_009731565,[27]) and ‘MEX_DaPlex’ (GCA_018135715,[39]). Note that chromosome numbers in the D_plex4 assembly are different, and chr15 corresponds to chr7. For alignment, we used both minimap2[40] with the ‘asm20’ parameter preset and the *nucmer* command in MUMmer v.4.0.0[41]. Alignments were explored visually using the online tool at https://dot.sandbox.bio.

For final visualisation, we used the minimap2 alignments to the outgroup *D. plexippus* ‘MEX_DaPlex’ assembly, and discarded alignments <500 bp in length. For comparison among assemblies of *D. chrysippus* supergene alleles, we used minimap2 with preset ‘asm10’, and discarded alignments of <2 kb and with divergence (‘dv value output by minimap2) <0.2.

To identify syntenic blocks and delineate rearranged regions, we developed an algorithm that merges adjacent alignments, implemented in the script asynt.R available at https://github.com/simonhmartin/asynt. The algorithm has three steps. First, alignments are split into ‘sub-blocks’ that each correspond to a unique tract of the reference assembly (each alignment corresponds to one sub-block, except overlapping intervals are defined as distinct sub-blocks). Second, sub-blocks below a minimum size in the reference are discarded. Third, adjacent sub-blocks that are in the same orientation and are below some threshold distance apart in the reference are merged to yield syntenic blocks. These three steps can be performed iteratively to first identify regions of fine-scale synteny and build these up into larger syntenic blocks. We used three iterations with minimum sub-block sizes of 200 bp, 2 kb and 20 kb, respectively, and a maximum distance between sub-blocks of 100 kb.

### Ancestry painting based on sequence divergence

We devised an approach to ‘paint’ each assembled haplotype according to its ancestry based on aligned short read data from representative individuals known to carry one of the three known supergene alleles. We first selected several representative wild individuals identified as homozygous for one of the three supergene alleles in a previous study[13] (see Table S1 for accession numbers). For each target assembly, short reads from the three sets of representative individuals were aligned using BWA mem v0.7.17 with default parameters. BAM files were generated using SAMtools v1.9 and sorted using PicardTools (https://broadinstitute.github.io/picard/) SortSam v2.21.1. Duplicate reads were identified and removed using PicardTools MarkDuplicates. Genotypes were called using the repeat-masked references with bcftools v1.10.2[29,42], using the mpileup, call and filter tools to retain only genotypes with an individual read depth (DP) ⩾8 and genotype quality (GQ) ⩾20. Diploid genotypes for each individual were exported, along with the haploid assembly genotype using the script parseVCF.py, and diversity (π) and divergence (*d*_*XY*_) between each reference panel and the assembly were computed using the script popgenWindows.py (https://github.com/simonhmartin/genomics_general). Non-overlapping windows of 25,000 genotyped sites were used, and for each reference panel, at least 10,000 sites had to be genotyped in at least one of the individuals. We then calculated *d*_*a*_ by subtracting π for a given reference panel from *d*_*XY*_. ‘Ancestry painting’ was performed by assigning, for each window, the reference panel that was most similar to the target assembly according to *d*_*a*_, with the added requirement that the difference between the lowest and second lowest *d*_*a*_ value had to be ⩾0.01 (Figure S2).

### Copy number variation

We estimated gene copy number variation based on read depth of Illumina reads. We used *D. plexippus* (Dplex_v4) as a reference as this species was inferred to represent the ancestral state in gene copy number. Reads were aligned and BAM files were processed as described above. To ensure that copy numbers were comparable among species and not biased by poor read mapping in highly divergent regions, we only considered read depth at exonic sites that had at least one read mapped in all individuals from all species. To this end, we generated a vcf file as described above, but without filtering for depth or genotype quality, and used bcftools view to retain only sites at which all individuals had non-zero depth. Finally, median depth was computed for each gene using the scripts parseVCF.py and windowStats.py (https://github.com/simonhmartin/genomics_general). Genes with <100 sites across all exons present in the data set were excluded. Gene copy numbers were estimated by normalising median depths by the median depth across all genes in the chromosome and rounding to the nearest whole number.

### Phylogenetic analyses

We generated alignments for the coding regions of each gene using available short-read Illumina sequence data for seven *Danaus* species and an outgroup from the sister genus *Tirumala* that was sequenced for this study (Table S1). The same VCF files described above (including invariant sites) for copy number analysis were used, with all genotypes with a genotype quality (GQ) <20 and read depth (DP) < 5 set to missing data. We then removed all sites with any missing data, converted diploid genotypes to single bases with heterozygous genotypes expressed as IUPAC ambiguity characters, and finally extracted alignments for each gene using the scripts parseVCF.py, filterGeno.py and extractCDSalignments.py available at https://github.com/simonhmartin/genomics_general. After generating these alignments for each gene, we discarded genes where alignments had >500 Ns and comprised ≥75% Ns.

For species tree inference, we further excluded all genes on chromosomes thought to carry rearrangements[13]: chr4 (chr11 in Dplex_v4), chr7 (chr15), chr15 (chr7), chr17 (chr16), chr22 (chr20), chr30 (chr28). The resulting 5954 gene alignments were analysed using the multispecies coalescent tool ASTRAL[43], implemented in the package ParGenes v.1.1.2 ([44]; which produces individual gene trees with RAxML[45] and then initiates an ASTRAL run on this full set of trees). Default parameters for ParGenes were used.

To explore the history of individual rearrangements we subsetted our filtered set of gene alignments to only the genes found within the bounds of the rearranged region (Region 1 was divided into two to account for the separate inversions in orientis and chrysippus). We then ran RAxML v.8.2.12[45] on a concatenated alignment for each subset using the GTRGAMMA model with 100 rapid bootstrap replicates (-f a -N 100).

### Sequence divergence in sliding windows

We calculated sequence divergence between reference panels representing each supergene allele using the same genotype data described in the ancestry painting section above. Non-overlapping windows of 25,000 genotyped sites were used, and ≥10,000 sites had to be genotyped in at least one of the individuals from each panel. Nucleotide diversity (π) and divergence (*d*_*XY*_) were computed using the script popgenWindows.py (https://github.com/simonhmartin/genomics_general), and we then calculated *d*_*a*_ by subtracting the average π for a given pair from *d*_*XY*_.

## DATA ACCESSIBILITY

All four assemblies and corresponding annotations are available at the European Nucleotide Archive project accession: XXXXX. Additional data files are provided at http://datadryad.org/XXX, including: repeat library, alignment files, genotype files, window-based divergence, repeat-content and read depth. Scripts for assembly polishing, the analysis of repeat content, genome annotation, and phylogenetic tree construction are available at https://github.com/RishiDeKayne/Danaus_supergene_structure. Scripts for genome alignment and synteny block inference, ancestry painting and divergence analyses, and read depth and copy number analyses are available at https://github.com/simonhmartin/Danaus_supergene_structure.

## ACKNOWLEDGEMENTS

We thank Alex Mackintosh for providing advice on genome assembly and annotation and valuable feedback on our interpretations of the results. We thank Chay Graham for helpful comments on the manuscript. This work was supported by a Royal Society University Research Fellowship (URF\R1\180682) and Enhancement Award (RGF\EA\181071) awarded to SHM, a Swiss National Science Foundation Early Postdoc Mobility Fellowship (P2BEP3_195567) awarded to RDK, and a National Geographic Society Research Grant (WW-138R-17) awarded to IJG.

## REFERENCES

1. Küpper C et al. 2016 A supergene determines highly divergent male reproductive morphs in the ruff. Nat. Genet. 48, 79–83.

2. Mérot C, Llaurens V, Normandeau E, Bernatchez L, Wellenreuther M. 2020 Balancing selection via life-history trade-offs maintains an inversion polymorphism in a seaweed fly. Nat. Commun. 11, 670.

3. Kirkpatrick M, Barton N. 2006 Chromosome inversions, local adaptation and speciation. Genetics 173, 419–434.

4. Wellenreuther M, Bernatchez L. 2018 Eco-evolutionary genomics of chromosomal inversions. Trends Ecol. Evol. 33, 427–440.

5. Joron M et al. 2011 Chromosomal rearrangements maintain a polymorphic supergene controlling butterfly mimicry. Nature 477, 203–206.

6. Charlesworth D, Charlesworth B, Marais G. 2005 Steps in the evolution of heteromorphic sex chromosomes. Heredity 95, 118–128.

7. Ozias-Akins P, Akiyama Y, Hanna WW. 2003 Molecular characterization of the genomic region linked with apomixis in Pennisetum/Cenchrus. Funct. Integr. Genomics 3, 94–104.

8. Li J et al. 2016 Genetic architecture and evolution of the S locus supergene in Primula vulgaris. Nat Plants 2, 16188.

9. Iijima T, Kajitani R, Komata S, Lin C-P, Sota T, Itoh T, Fujiwara H. 2018 Parallel evolution of Batesian mimicry supergene in two Papilio butterflies, P. polytes and P. memnon. Sci Adv 4, eaao5416.

10. Yan Z et al. 2020 Evolution of a supergene that regulates a trans-species social polymorphism. Nat Ecol Evol 4, 240–249.

11. Brower LP, Edmunds M, Moffitt CM. 1975 Cardenolide content and palatability of a population of Danaus chrysippus butterflies from West Africa. Journal of Entomology Series A, General Entomology 49, 183–196.

12. Smith DAS, Owen DF, Gordon IJ, Lowis NK. 1997 The butterfly Danaus chrysippus (L.) in East Africa: polymorphism and morph-ratio clines within a complex, extensive and dynamic hybrid zone. Zool. J. Linn. Soc. 120, 51–78.

13. Martin SH et al. 2020 Whole-chromosome hitchhiking driven by a male-killing endosymbiont. PLoS Biol. 18, e3000610.

14. Smith DAS. 1979 The significance of beak marks on the wings of an aposematic, distasteful and polymorphic butterfly. Nature 281, 215–216.

15. Smith DAS, Gordon IJ, Allen JA. 2010 Reinforcement in hybrids among once isolated semispecies of Danaus chrysippus(L.) and evidence for sex chromosome evolution. Ecol. Entomol. 35, 77–89.

16. Koren S et al. 2018 De novo assembly of haplotype-resolved genomes with trio binning. Nat. Biotechnol. 36, 1174–1182.

17. Singh KS, De-Kayne R, Omufwoko KS, Martins D, Bass C, ffrench-Constant R, Martin SH. 2021 Genome assembly of Danaus chrysippus and comparison with the Monarch Danaus plexippus. Biorxiv, BIORXIV/2021/470194.

18. Chazot N et al. 2019 Priors and Posteriors in Bayesian Timing of Divergence Analyses: The Age of Butterflies Revisited. Syst. Biol. 68, 797–813.

19. Cruickshank TE, Hahn MW. 2014 Reanalysis suggests that genomic islands of speciation are due to reduced diversity, not reduced gene flow. Mol. Ecol. 23, 3133–3157.

20. Montgomery EA, Huang SM, Langley CH, Judd BH. 1991 Chromosome rearrangement by ectopic recombination in Drosophila melanogaster: genome structure and evolution. Genetics 129, 1085–1098.

21. Arguello JR, Connallon T. 2011 Gene duplication and ectopic gene conversion in Drosophila. Genes 2, 131–151.

22. Puerma E, Orengo DJ, Aguadé M. 2016 Multiple and diverse structural changes affect the breakpoint regions of polymorphic inversions across the Drosophila genus. Sci. Rep. 6, 36248.

23. Maggiolini FAM et al. 2020 Single-cell strand sequencing of a macaque genome reveals multiple nested inversions and breakpoint reuse during primate evolution. Genome Res. 30, 1680–1693.

24. Corbett-Detig RB, Said I, Calzetta M, Genetti M, McBroome J, Maurer NW, Petrarca V, Della Torre A, Besansky NJ. 2019 Fine-Mapping Complex Inversion Breakpoints and Investigating Somatic Pairing in the Anopheles gambiae Species Complex Using Proximity-Ligation Sequencing. Genetics 213, 1495–1511.

25. Otto SP et al. 2011 About PAR: the distinct evolutionary dynamics of the pseudoautosomal region. Trends Genet. 27, 358–367.

26. Korunes KL, Noor MAF. 2019 Pervasive gene conversion in chromosomal inversion heterozygotes. Mol. Ecol. 28, 1302–1315.

27. Gu L, Reilly PF, Lewis JJ, Reed RD, Andolfatto P, Walters JR. 2019 Dichotomy of Dosage Compensation along the Neo Z Chromosome of the Monarch Butterfly. Curr. Biol. 29, 4071–4077.e3.

28. Li H, Durbin R. 2010 Fast and accurate long-read alignment with Burrows-Wheeler transform. Bioinformatics 26, 589–595.

29. Li H et al. 2009 The Sequence Alignment/Map format and SAMtools. Bioinformatics 25, 2078–2079.

30. Barnett DW, Garrison EK, Quinlan AR, Strömberg MP, Marth GT. 2011 BamTools: a C++ API and toolkit for analyzing and managing BAM files. Bioinformatics 27, 1691–1692.

31. Waples RK, Albrechtsen A, Moltke I. 2019 Allele frequency-free inference of close familial relationships from genotypes or low-depth sequencing data. Mol. Ecol. 28, 35–48.

32. Koren S, Walenz BP, Berlin K, Miller JR, Bergman NH, Phillippy AM. 2017 Canu: scalable and accurate long-read assembly via adaptive k-mer weighting and repeat separation. Genome Res. 27, 722–736.

33. Cheng H, Concepcion GT, Feng X, Zhang H, Li H. 2021 Haplotype-resolved de novo assembly using phased assembly graphs with hifiasm. Nat. Methods 18, 170–175.

34. Nurk S et al. 2020 HiCanu: accurate assembly of segmental duplications, satellites, and allelic variants from high-fidelity long reads. Genome Res. (doi:10.1101/gr.263566.120)

35. Simão FA, Waterhouse RM, Ioannidis P, Kriventseva EV, Zdobnov EM. 2015 BUSCO: assessing genome assembly and annotation completeness with single-copy orthologs. Bioinformatics 31, 3210–3212.

36. Roach MJ, Schmidt SA, Borneman AR. 2018 Purge Haplotigs: allelic contig reassignment for third-gen diploid genome assemblies. BMC Bioinformatics 19, 460.

37. Smit AFA, Hubley R, Green P. 2015 RepeatMasker Open-4.0. 2013--2015.

38. Hoff KJ, Lomsadze A, Stanke M, Borodovsky M. 2018 BRAKER2: incorporating protein homology information into gene prediction with GeneMark-EP and AUGUSTUS. Plant and Animal Genomes XXVI

39. Ranz JM et al. 2021 A de novo transcriptional atlas in Danaus plexippus reveals variability in dosage compensation across tissues. Commun Biol 4, 791.

40. Li H. 2018 Minimap2: pairwise alignment for nucleotide sequences. Bioinformatics 34, 3094–3100.

41. Marçais G, Delcher AL, Phillippy AM, Coston R, Salzberg SL, Zimin A. 2018 MUMmer4: A fast and versatile genome alignment system. PLoS Comput. Biol. 14, e1005944.

42. Danecek P et al. 2021 Twelve years of SAMtools and BCFtools. Gigascience 10. (doi:10.1093/gigascience/giab008)

43. Mirarab S, Reaz R, Bayzid MS, Zimmermann T, Swenson MS, Warnow T. 2014 ASTRAL: genome-scale coalescent-based species tree estimation. Bioinformatics 30, i541–8.

44. Morel B, Kozlov AM, Stamatakis A. 2019 ParGenes: a tool for massively parallel model selection and phylogenetic tree inference on thousands of genes. Bioinformatics 35, 1771–1773.

45. Stamatakis A. 2014 RAxML version 8: a tool for phylogenetic analysis and post-analysis of large phylogenies. Bioinformatics 30, 1312–1313.

