## Supplementary material for "Stepwise evolution of a butterfly supergene via duplication and inversion": Tables S2-S4 and Figures S1-S11

**Table S1.** Sample data and accession numbers

Separate .xlsx file

**Table S2**. Summary of Trio-binning for PacBio sequences

| Broods | MB18102 | | SB211* | |
|---|---|---|---|---|
| | reads | bp | reads | bp |
| raw reads | 2,737,624 | 34,667,556,950 | 1,432,815 | 12,524,291,964 |
| binned to mother | 1,108,254 | 16,195,119,264 | 729,759 | 6,417,342,625 |
| binned to father | 1,197,570 | 18,042,353,099 | 685,070 | 6,051,324,571 |
| binned to unknown | 1,493 | 6,635,278 | 6,603 | 44,297,006 |
| too short | 430,307 | 423,449,309 | 11,383 | 11,327,762 |

*pooled reads of three offspring (see Text for more detail).

**Table S3**. Summary of assembly for Trio-binned PacBio reads.

| haplotype | MB18102MAT | MB18102PAT | SB211MAT | SB211PAT | SB211MAT | SB211PAT |
|---|---|---|---|---|---|---|
| **Primary assembly** | | | | | | |
| Trio-binning | canu | canu | canu | canu | canu | canu |
| assembly | canu | canu | HiCanu | HiCanu | hifiasm | hifiasm |
| seqs | 2,322 | 2,432 | 5,492 | 4,134 | 3,010 | 2,131 |
| size | 413,742,804 | 502,480,781 | 623,429,130 | 571,050,471 | 605,714,309 | 539,605,120 |
| N50 | 609,454 | 3,330,071 | 264,060 | 517,254 | 548,905 | 1,068,083 |
| min | 2,040 | 5,413 | 4,780 | 4,055 | 8,488 | 8,676 |
| max | 12,670,935 | 17,388,644 | 16,188,425 | 9,794,742 | 13,066,125 | 13,931,720 |
| N75 | 171,894 | 193,046 | 99,734 | 140,811 | 193,782 | 307,062 |
| N90 | 58,142 | 59,311 | 43,510 | 51,312 | 79,870 | 94,420 |
| auN | 3,490,769.02 | 4,529,070.53 | 2,019,462.67 | 1,916,794.27 | 2,136,138.64 | 2,662,837.03 |
| **Cleaned assembly*** | | | | | | |
| seqs | 722 | 208 | 946 | 751 | 517 | 394 |
| size | 280,793,503 | 318,388,826 | 321,290,812 | 331,935,336 | 320,813,483 | 330,120,261 |
| N50 | 3,045,643 | 7,070,092 | 961,241 | 2,065,040 | 2,103,362 | 2,813,076 |
| min | 2,653 | 8,881 | 7,512 | 5,515 | 9,282 | 10,912 |
| max | 12,670,935 | 17,388,644 | 16,188,425 | 9,794,742 | 13,066,125 | 13,931,720 |
| N75 | 353,289 | 3,639,342 | 308,110 | 456,385 | 673,485 | 1,003,716 |
| N90 | 145,879 | 1,107,980 | 156,280 | 201,470 | 307,963 | 461,361 |
| auN | 5,002,700.16 | 6,981,774.17 | 3,700,651.77 | 3,032,044.92 | 3,741,985.28 | 3,994,391.24 |
| further analysis | yes | yes | yes | no | no | yes |

*cleaned by using Purge Haplotigs. All statistics were generated by N50 (https://metacpan.org/pod/Proch::N50)

**Table S4**. Summary of BUSCO analysis.

| **Assembly** | **C** | **S** | **D** | **F** | **M** | **n** |
|---|---|---|---|---|---|---|
| *arthropoda_odb9* | | | | | | |
| MB18102PAT | 1,055 (99.0%) | 919 (86.2%) | 136 (12.8%) | 4 (0.4%) | 7 (0.6%) | 1,066 |
| MB18102MAT | 968 (90.8%) | 958 (89.9%) | 10 (0.9%) | 15 (1.4%) | 83 (7.8%) | 1,066 |
| SB211PAT | 1,053 (98.8%) | 1,020 (95.7%) | 33 (3.1%) | 3 (0.3%) | 10 (0.9%) | 1,066 |
| SB211MAT | 992 (93.1%) | 937 (87.9%) | 55 (5.2%) | 3 (0.3%) | 71 (6.6%) | 1,066 |
| *insecta_odb9* | | | | | | |
| MB18102PAT | 1,628 (98.2%) | 1,414 (85.3%) | 214 (12.9%) | 11 (0.7%) | 19 (1.1%) | 1,658 |
| MB18102MAT | 1,482 (89.4%) | 1,461 (88.1%) | 21 (1.3%) | 32 (1.9%) | 144 (8.7%) | 1,658 |
| SB211PAT | 1,625 (98.0%) | 1,577 (95.1%) | 48 (2.9%) | 7 (0.4%) | 26 (1.6%) | 1,658 |
| SB211MAT | 1,536 (92.7%) | 1,462 (88.2%) | 74 (4.5%) | 9 (0.5%) | 113 (6.8%) | 1,658 |

**C**, complete; **S,** complete and single-copy; **D**, complete and duplicated; **F**, fragmented; **M**, missing; **n**, total BUSCO groups searched

**Table S5.** Gene copy number estimates for high-copy Region 3 genes
Separate .xlsx file.

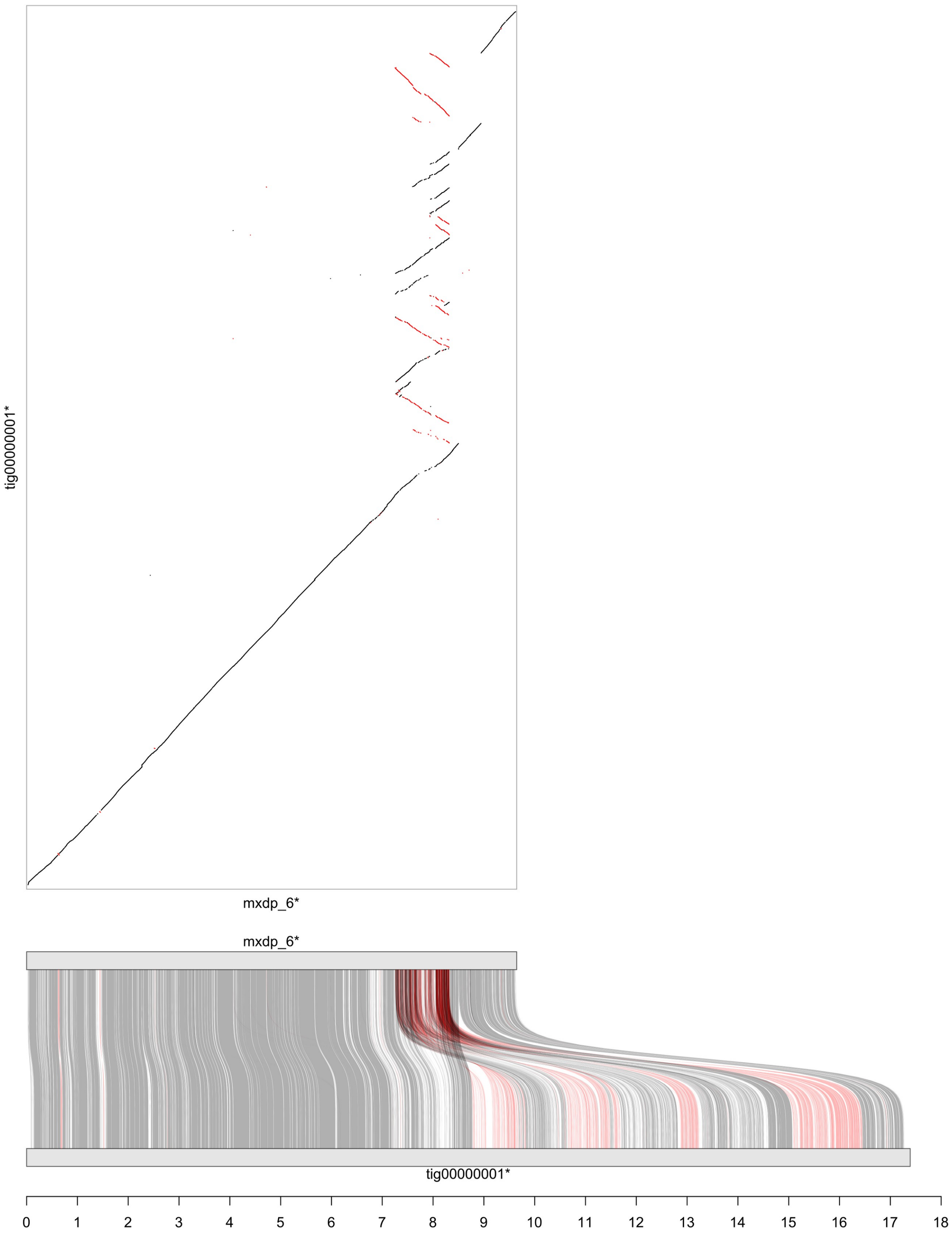


**Figure S1A.** Alignment of *D. chrysippus* (MB18102PAT assembly) to *D. plexippus* (MEX_DaPlex assembly) for chromosome 15. Alignment was performed using minimap2 (see Methods). Only alignments of at least 500 bp in length are shown, with grey indicating forward orientation and red indicating reverse orientation. Contig names followed by an asterisk indicates that the contig was reversed for improved visualisation. The x-axis is scaled in megabases.

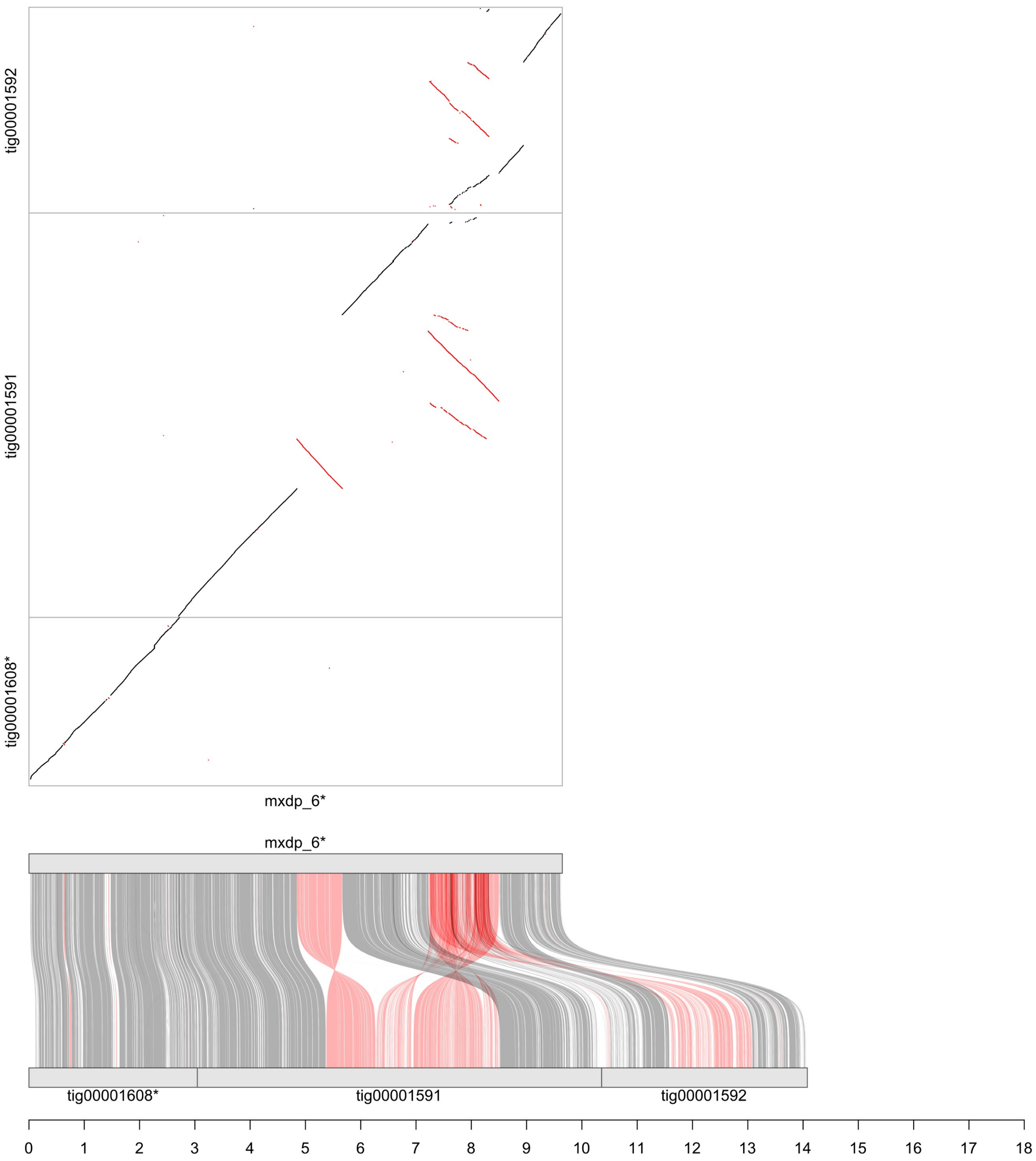


**Figure S1B.** Alignment of *D. chrysippus* (MB18102MAT assembly) to *D. plexippus* (MEX_DaPlex assembly) for chromosome 15. Alignment was performed using minimap2 (see Methods). Only alignments of at least 500 bp in length are shown, with grey indicating forward orientation and red indicating reverse orientation. Contig names followed by an asterisk indicates that the contig was reversed for improved visualisation. The x-axis is scaled in megabases.

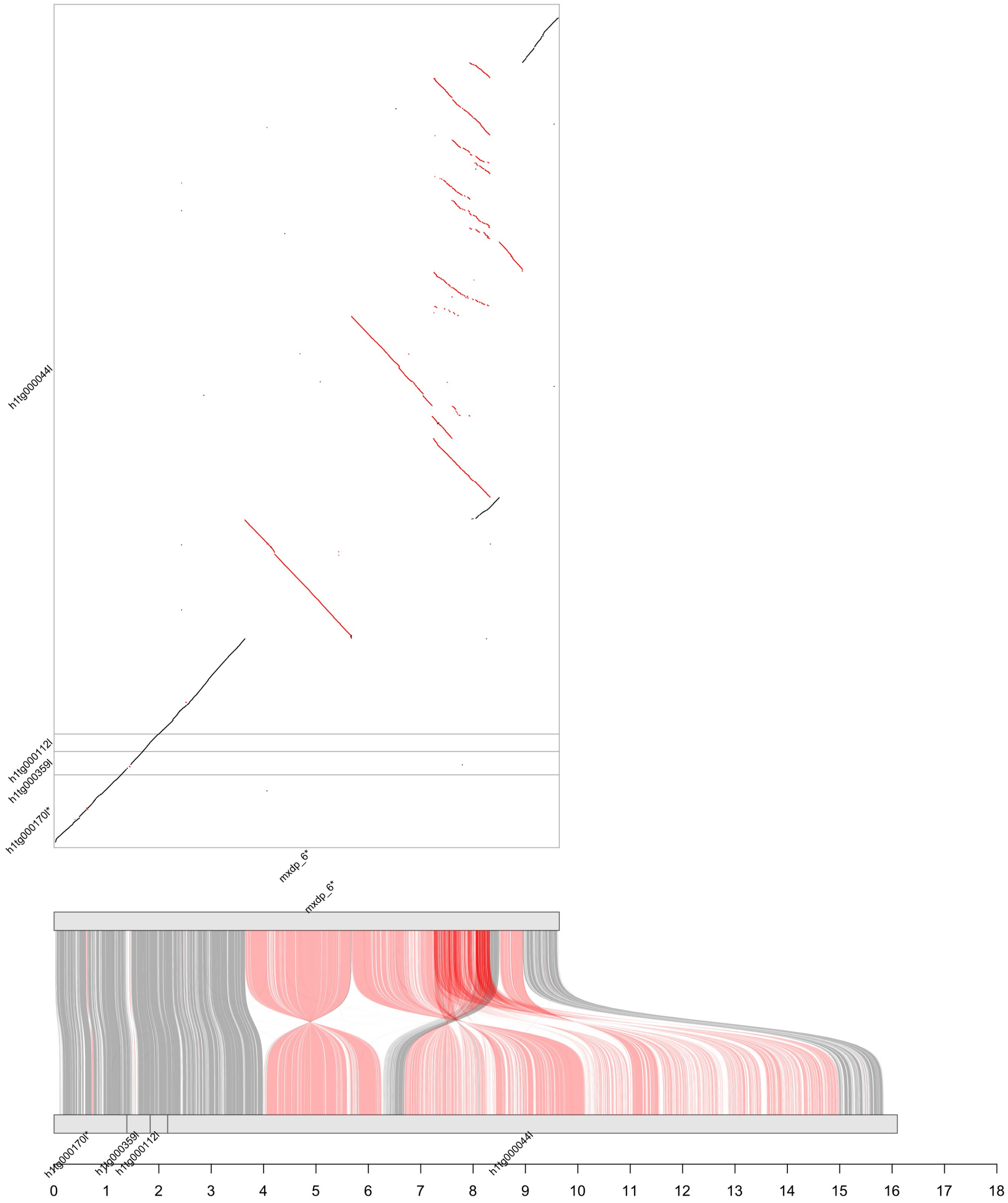


**Figure S1C.** Alignment of *D. chrysippus* (SB211PAT assembly) to *D. plexippus* (MEX_DaPlex assembly) for chromosome 15. Alignment was performed using minimap2 (see Methods). Only alignments of at least 500 bp in length are shown, with grey indicating forward orientation and red indicating reverse orientation. Contig names followed by an asterisk indicates that the contig was reversed for improved visualisation. The x-axis is scaled in megabases.

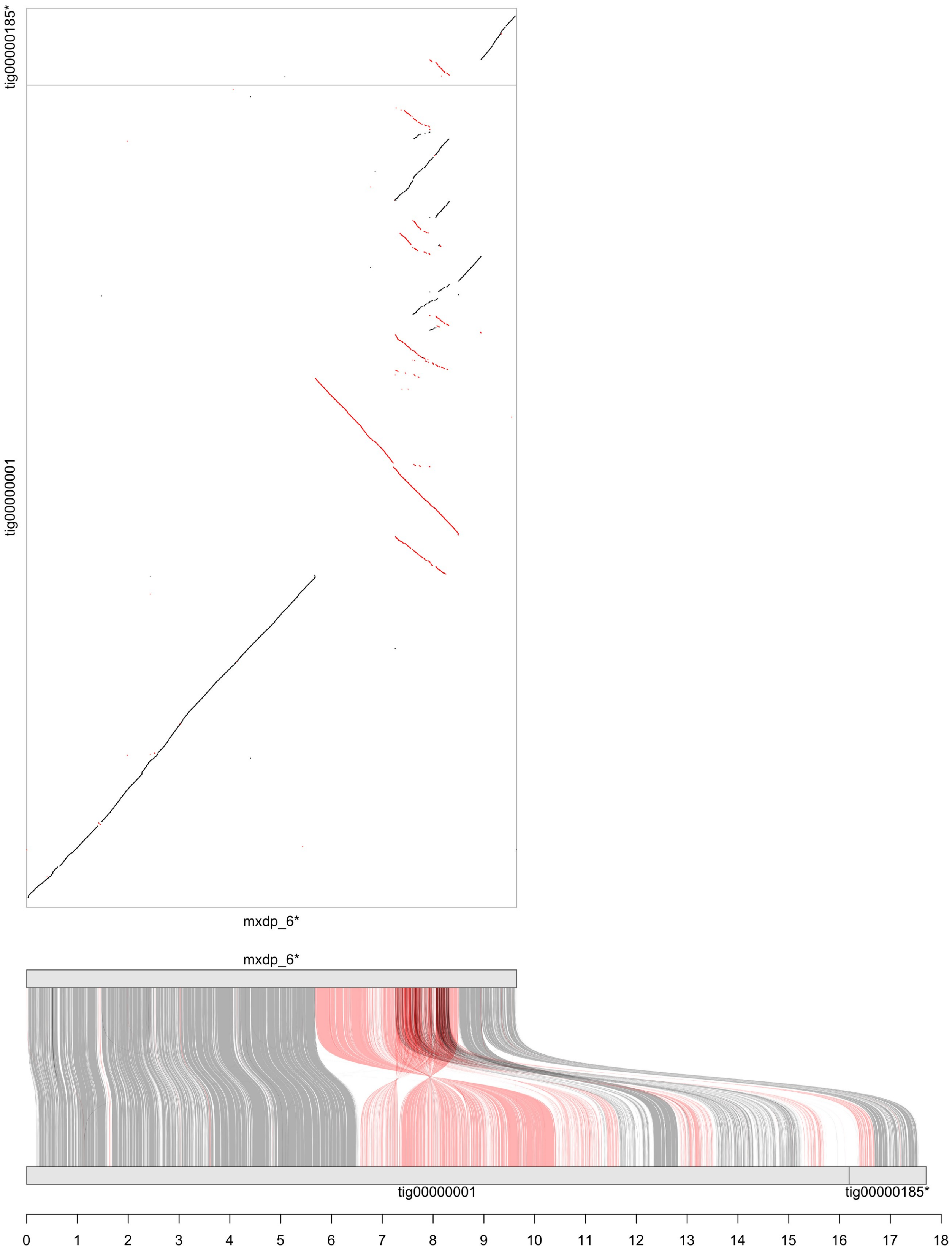


**Figure S1D.** Alignment of *D. chrysippus* (SB211MAT assembly) to *D. plexippus* (MEX_DaPlex assembly) for chromosome 15. Alignment was performed using minimap2 (see Methods). Only alignments of at least 500 bp in length are shown, with grey indicating forward orientation and red indicating reverse orientation. Contig names followed by an asterisk indicates that the contig was reversed for improved visualisation. The x-axis is scaled in megabases.

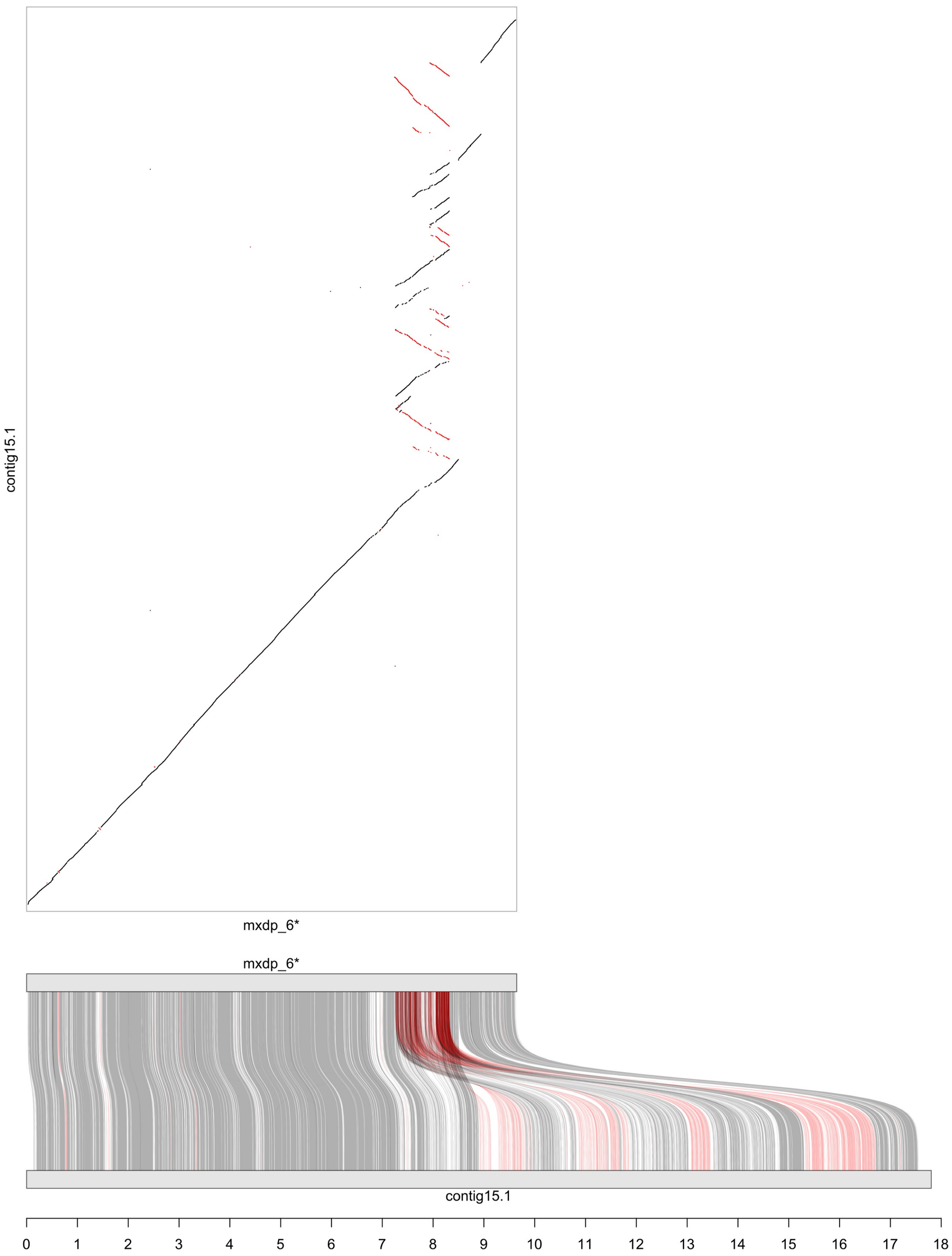


**Figure S1E.** Alignment of *D. chrysippus* (Dchry2.2 assembly) to *D. plexippus* (MEX_DaPlex assembly) for chromosome 15. Alignment was performed using minimap2 (see Methods). Only alignments of at least 500 bp in length are shown, with grey indicating forward orientation and red indicating reverse orientation. Contig names followed by an asterisk indicates that the contig was reversed for improved visualisation. The x-axis is scaled in megabases.

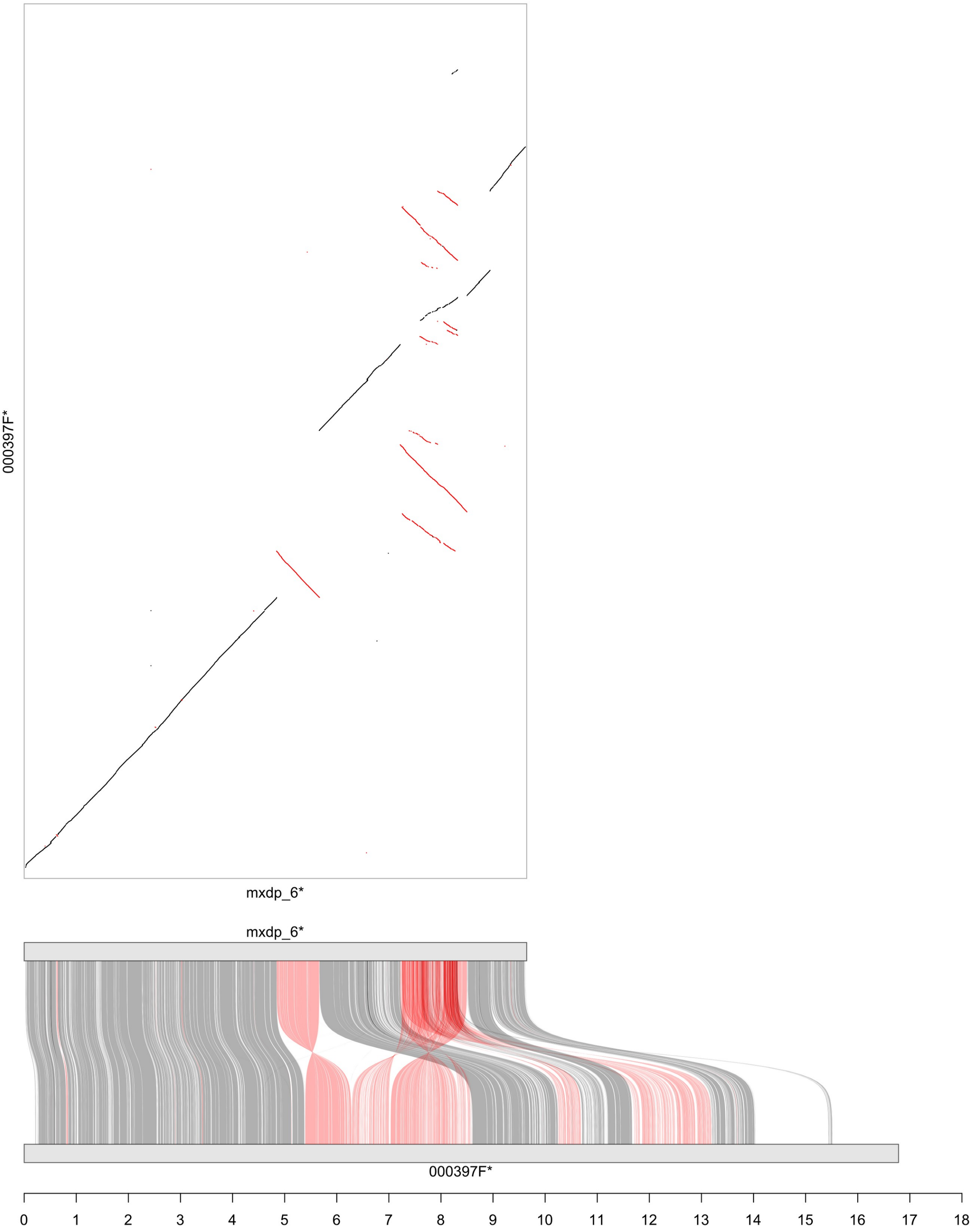


**Figure S1F.** Alignment of *D. chrysippus* (Dchry2 assembly alternative haplotype) to *D. plexippus* (MEX_DaPlex assembly) for chromosome 15. Alignment was performed using minimap2 (see Methods). Only alignments of at least 500 bp in length are shown, with grey indicating forward orientation and red indicating reverse orientation. Contig names followed by an asterisk indicates that the contig was reversed for improved visualisation. The x-axis is scaled in megabases. The additional fragment on the right-hand end is suspected to be telomeric repeats.

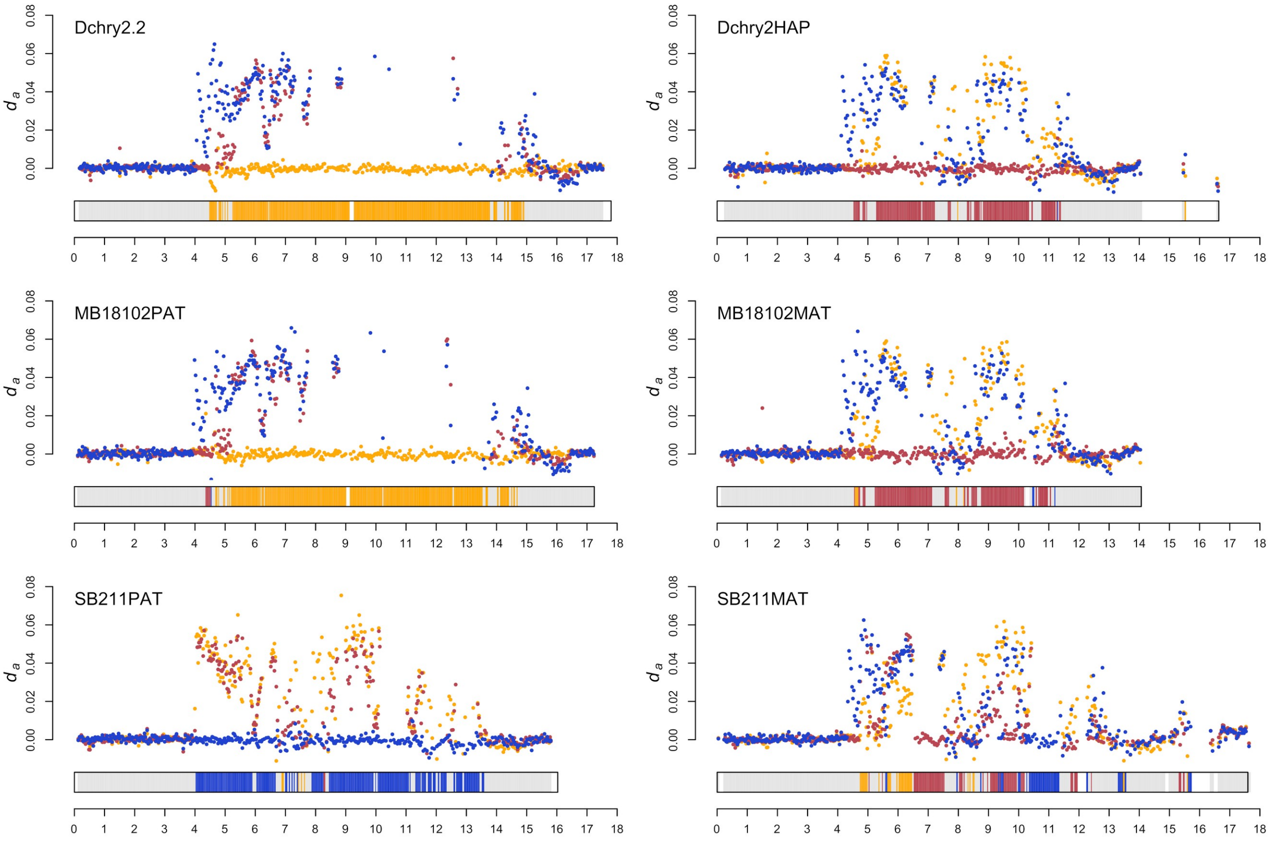


**Figure S2**. Ancestry painting identifies the chromosome 15 haplotype represented by each assembly. Lines represent *da* (absolute sequence divergence corrected for within-population diversity) between the haploid assembly and a panel of wild reference individuals that are homozygous for each of the three alleles: klugii (yellow), chrysippus (red) and orientis (blue) in windows of 25,000 genotyped sites. For each window, ancestry was assigned to the source population with the lowest divergence from the assembly sequence. Ancestry assignment is indicated by coloured blocks in the chromosome at the base of each plot. Windows in which no clear ancestry could be assigned (when the difference in *da* between the lowest and next-lowest is less than 0.01) are shown in grey. Small number of blocks are white, indicating stretches of ≥100,000 sites in which ≤25,000 sites could be genotyped. This is the case for a large portion on the right-hand end of the Dchry2 alternative haplotype assembly (Dchy2HAP, top right), which may be telomeric repeats.

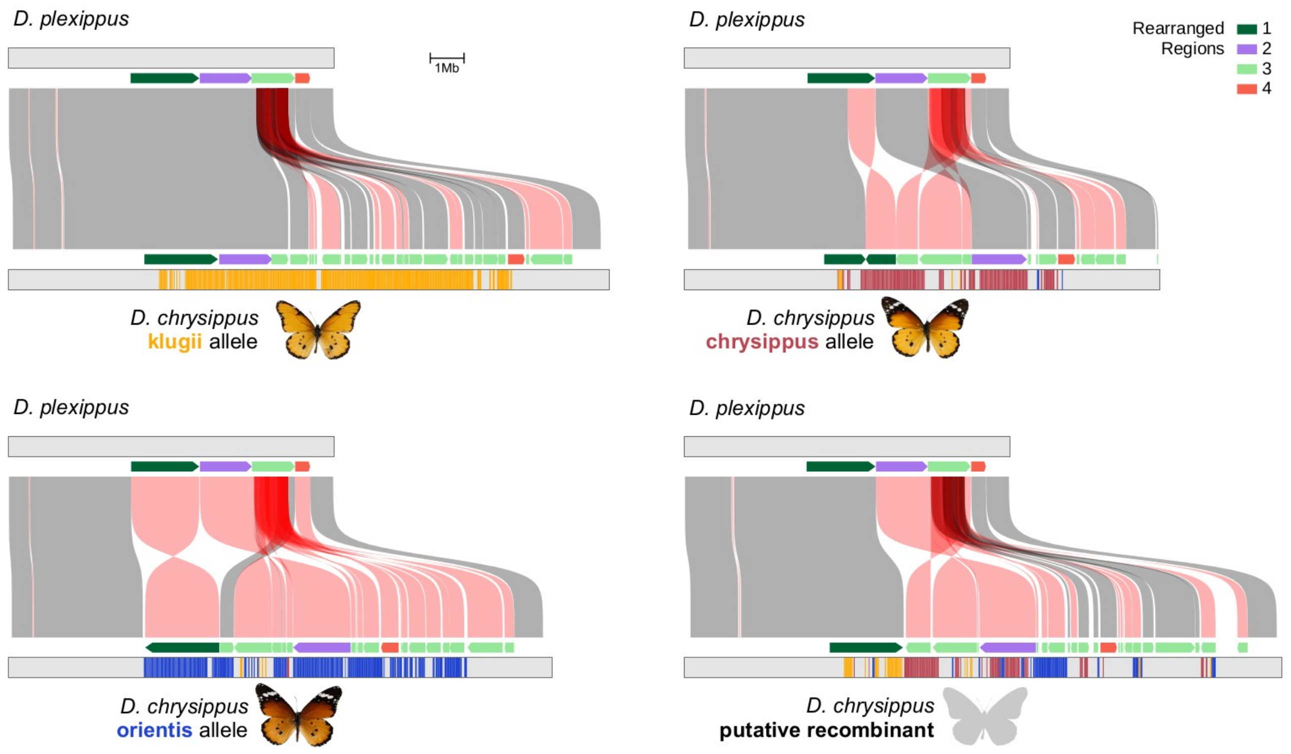


**Figure S3.** Synteny between the outgroup (*D. plexippus*) and each of the four assembled supergene alleles in *D. chrysippus*. Connecting boxes indicate syntenic blocks (see Methods for details) that are either in the forward (grey) or reverse (red) orientation. Boxes are partially transparent to show regions that are duplicated. Coloured arrows indicate the four regions that we identified as having experienced distinct rearrangements in *D. chrysippus*. The coloured bars indicate 'ancestry painting' (see Methods for details) in 50 kb windows along the *D. chrysippus* chromosomes. Regions lacking coloured bars could not be assigned ancestry because they showed sequence similarity to two or more different morphs.

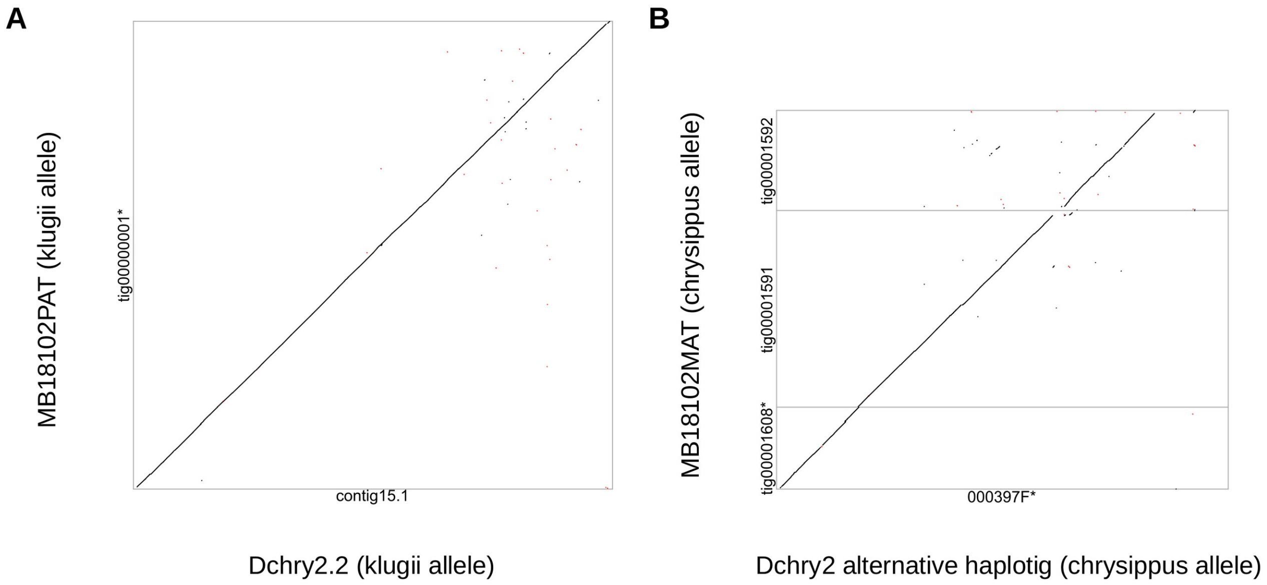


**Figure S4.** Alignment of independent assemblies of the same alleles to confirm assembly correctness. **A.** New assembly MB18102PAT is compared with previously-published assembly Dchry2.2, both of which carry the klugii allele. **B.** New assembly MB18102MAT is compared with the alternative haplotig for chr15 from the Dchry2 assembly, both of which represent the chrysippus allele. Alignment was performed using minimap2 (see Methods). Only alignments of at least 2000 bp and sequence divergence < 20% are shown, with grey indicating forward orientation and red indicating reverse orientation. Contig names followed by an asterisk indicates that the contig was reversed for improved visualisation. The additional portion of the Dchry2 alternative haplotype assembly (right) is suspected to be telomeric repeats (Figure S2).

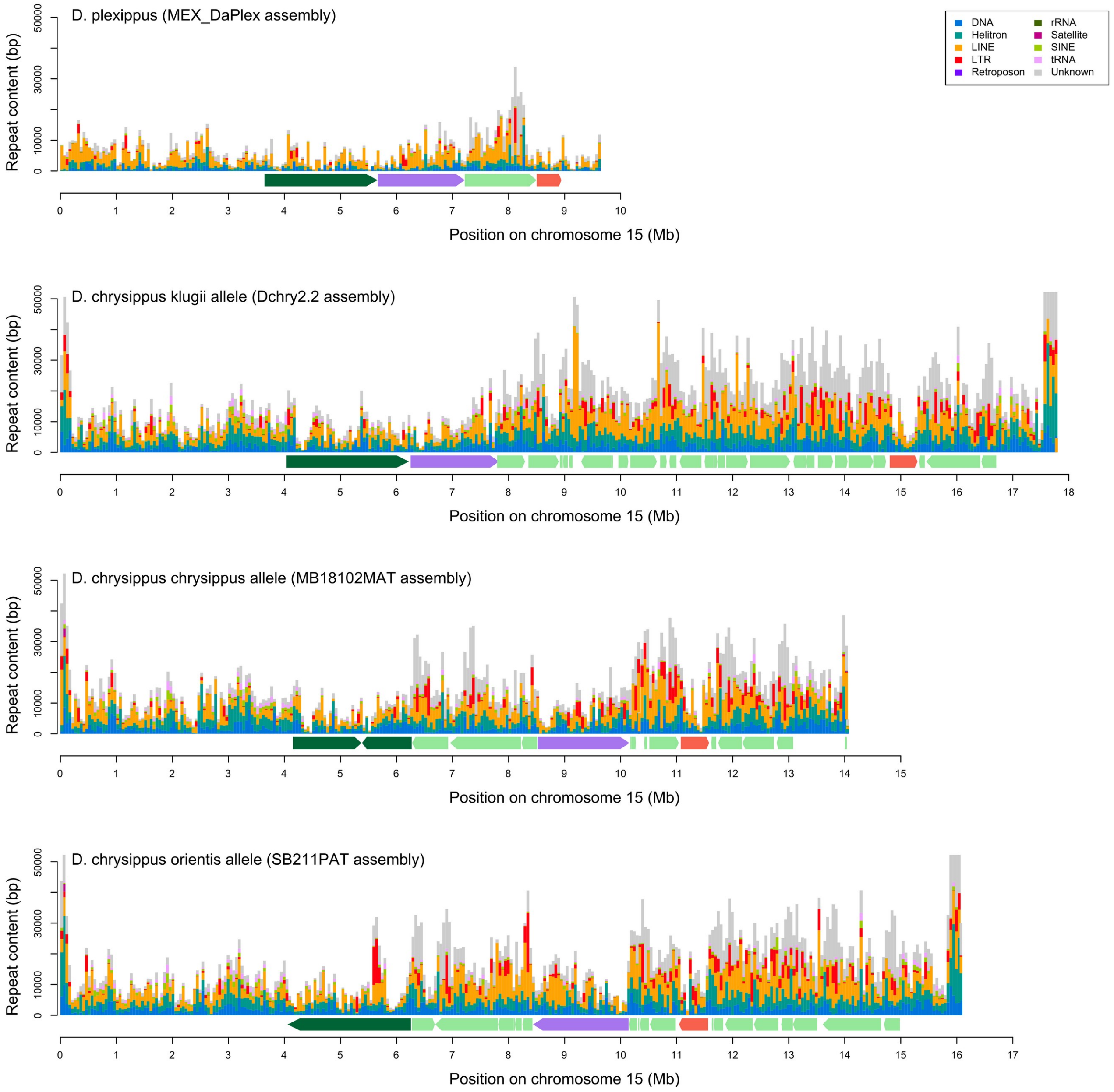


**Figure S5**. Repeat content on chromosome 15. Repeat content is plotted in 50 kb windows with different repeat types indicated by different colours. Block arrows along the bottom of each plot indicate the four distinct rearranged regions shown in Figure 1.

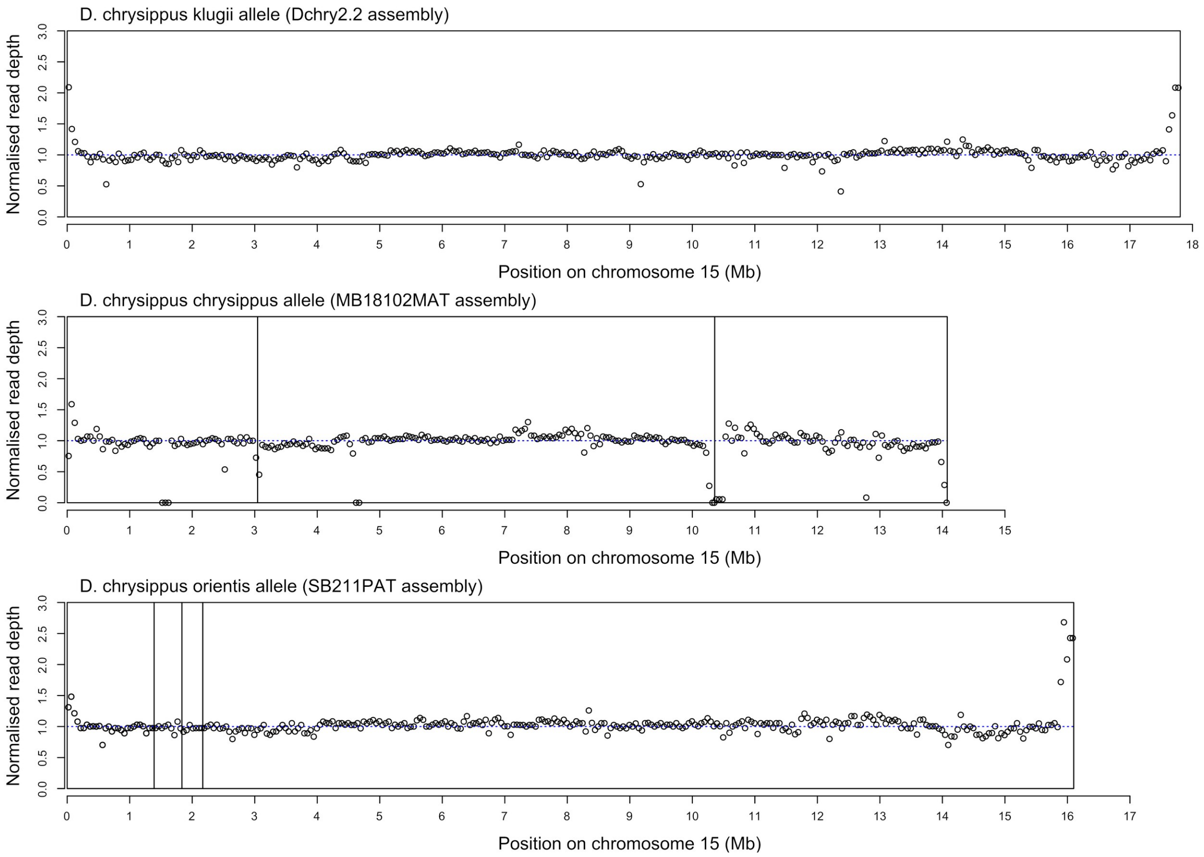


**Figure S6**. Plots of normalised read depth to test for collapsed repeats. Points represent the normalised median read depth in 50 kb windows for short reads mapped to each assembly, averaged over 2-4 individuals. For each allele, wild representative individuals known to be homozygous for the allele were used (Table S1). Boxes represent assembly contigs that have been ordered and oriented (see Figure S1). A normalised read depth of 1 (dashed line) is expected if the assembly is accurate. The lack of values approaching 2 or more indicates that there are few large collapsed repeats in our assembly, with the exception of the chromosome ends, which likely contain telomeric repeats.

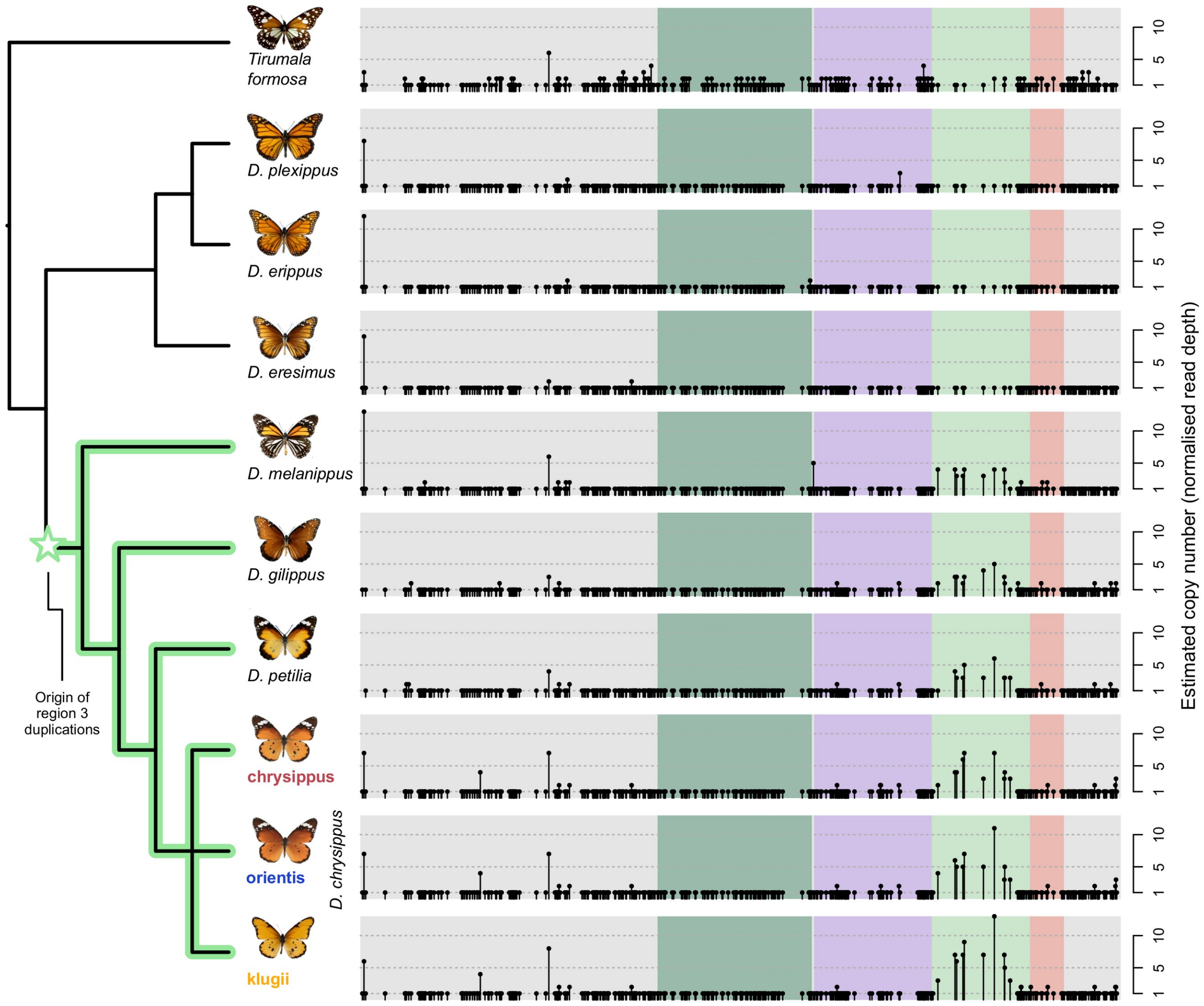


**Figure S7.** Copy number variation shows a deep origin of duplications on chromosome 15. For each species (or *D. chrysippus* morph), vertical black lines indicate the estimated copy number for each gene on chr15. Genes are positioned according to the *Danaus plexippus* (Dplex_v4 assembly). The four bands of coloured shading indicate Regions 1, 2, 3 and 4 as indicated in Figure 1. For species in which multiple individuals were available, copy number estimates were averaged (see Table S5 for individual copy numbers). The uneven copy number estimates in the outgroup *Tirumala formosa* probably largely reflect poor alignment from this diverged taxon to the *D. plexippus* reference. The tree on the left represents the species tree inferred using ASTRAL. All species in the clade shaded in green showed elevated copy numbers for ten genes in Region 3 of the chromosome, and the star indicates the assumed origin of the Region 3 duplications. Attributions for images not by the authors:

"MCZ:Ent:165651 Tirumala formosa formosa" - Tirumala formosa collected in Kenya by Museum of Comparative Zoology, Harvard University (licensed under http://creativecommons.org/licenses/by-nc-sa/3.0/)

"MCZ:Ent:152519 Danaus plexippus" - Danaus plexippus (Linnaeus, 1758) collected in United States of America by Museum of Comparative Zoology, Harvard University (licensed under http://creativecommons.org/licenses/by-nc-sa/3.0/)

"MCZ:Ent:164619 Danaus erippus" - Danaus erippus Cramer, 1775 collected in Brazil by Museum of Comparative Zoology, Harvard University (licensed under http://creativecommons.org/licenses/by-nc-sa/3.0/)

"MCZ:Ent:152526 Danaus eresimus tethys" - Danaus (Anosia) eresimus subsp. tethys Forbes, 1943 collected in United States of America by Museum of Comparative Zoology, Harvard University (licensed under http://creativecommons.org/licenses/by-nc-sa/3.0/)

"MCZ:Ent:210362 Danaus melanippus" - Danaus (Anosia) melanippus Cramer, 1777 collected in Thailand by Museum of Comparative Zoology, Harvard University (licensed under http://creativecommons.org/licenses/by-nc-sa/3.0/)

"MCZ:Ent:152524 Danaus gilippus berenice" - Danaus (Anosia) gilippus subsp. berenice Cramer, 1779 collected in United States of America by Museum of Comparative Zoology, Harvard University (licensed under http://creativecommons.org/licenses/by-nc-sa/3.0/)

"MCZ:Ent:89405 Danaus petilia" - Danaus petilia Stoll, 1790 collected in Australia by Museum of Comparative Zoology, Harvard University (licensed under http://creativecommons.org/licenses/by-nc-sa/3.0/)

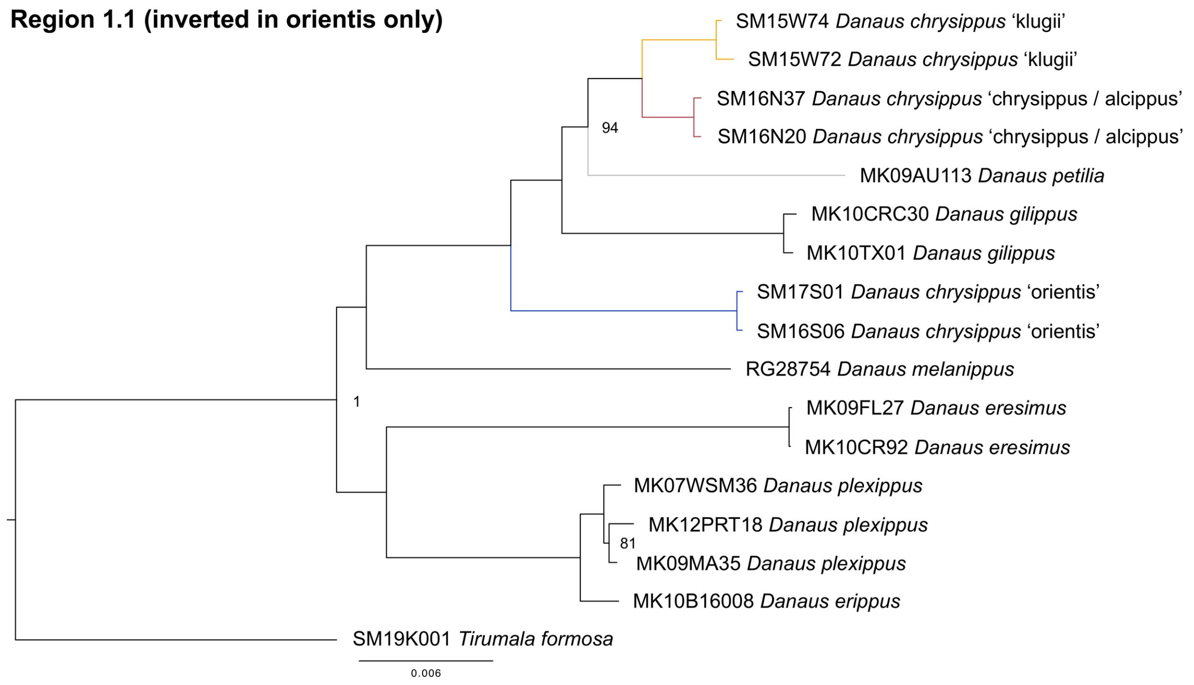


**Figure S8A.** Maximum likelihood phylogenies for concatenated coding sequences for Region 1.1 (inverted in orientis only; comprising 38 genes). All nodes had bootstrap support of 100% unless otherwise indicated, and nodes with <70% support are shown as polytomies.

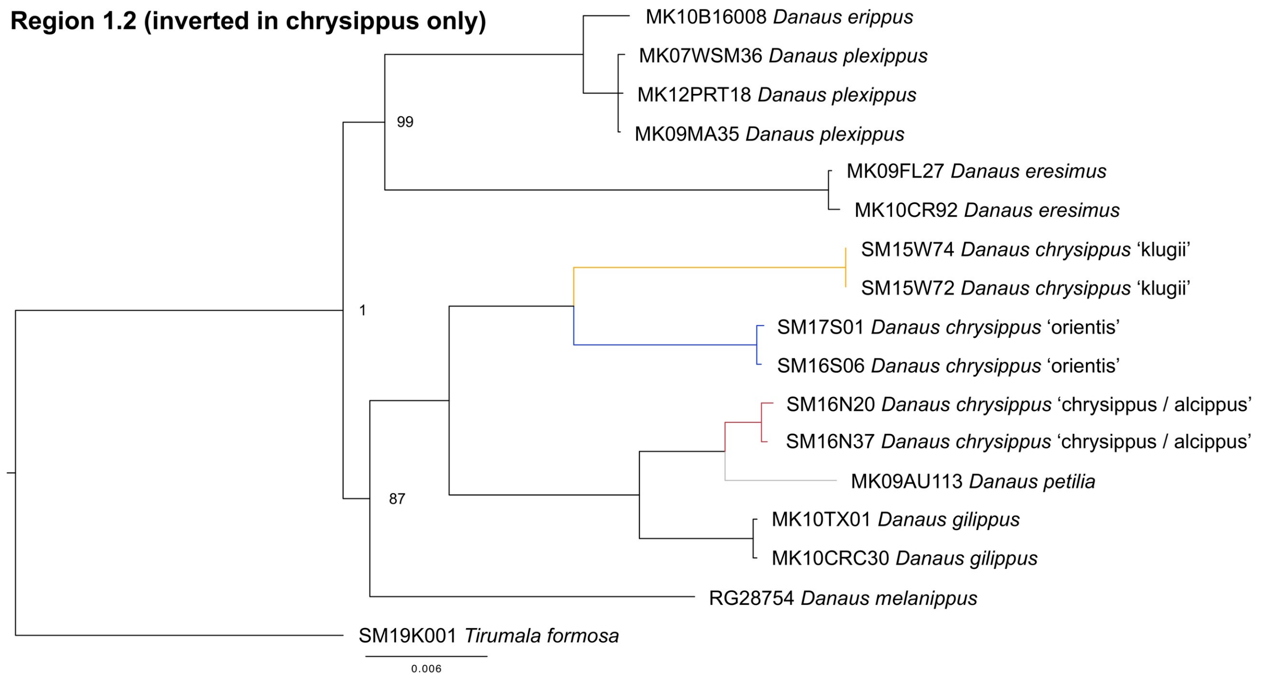


**Figure S8B.** Maximum likelihood phylogenies for concatenated coding sequences for Region 1.1 (inverted in orientis only; comprising 38 genes). All nodes had bootstrap support of 100% unless otherwise indicated, and nodes with <70% support are shown as polytomies.

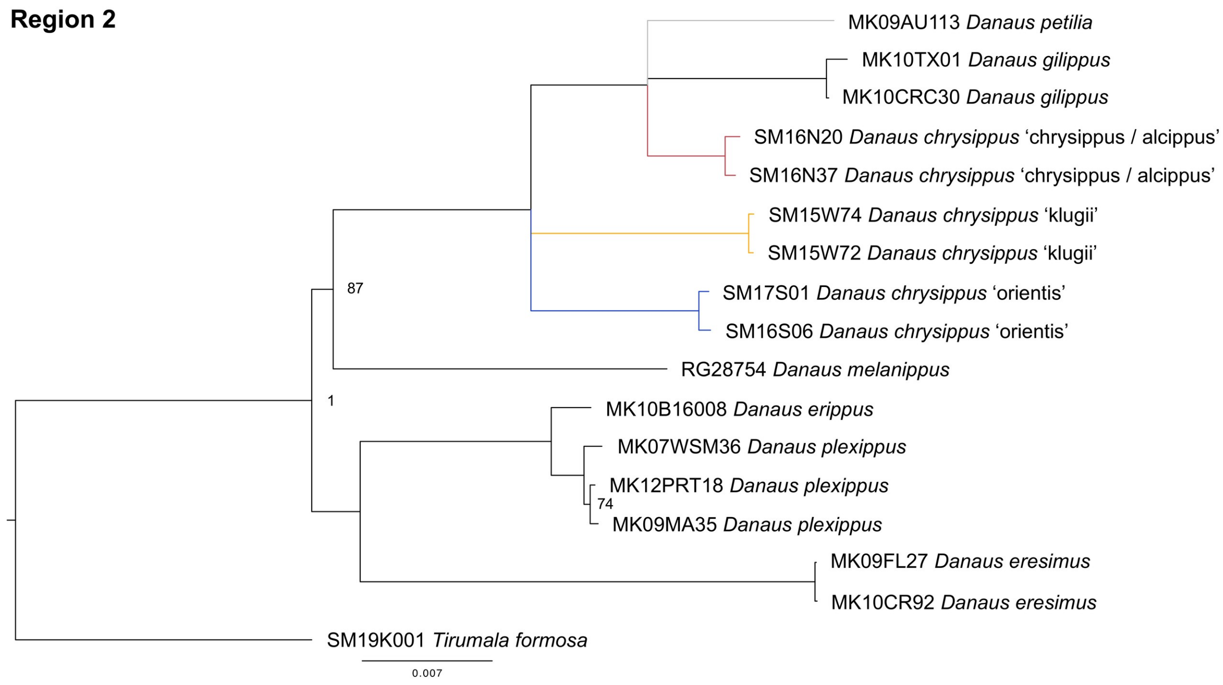


**Figure S8C.** Maximum likelihood phylogenies for concatenated coding sequences for Region 2 (comprising 61 genes). All nodes had bootstrap support of 100% unless otherwise indicated, and nodes with <70% support are shown as polytomies.

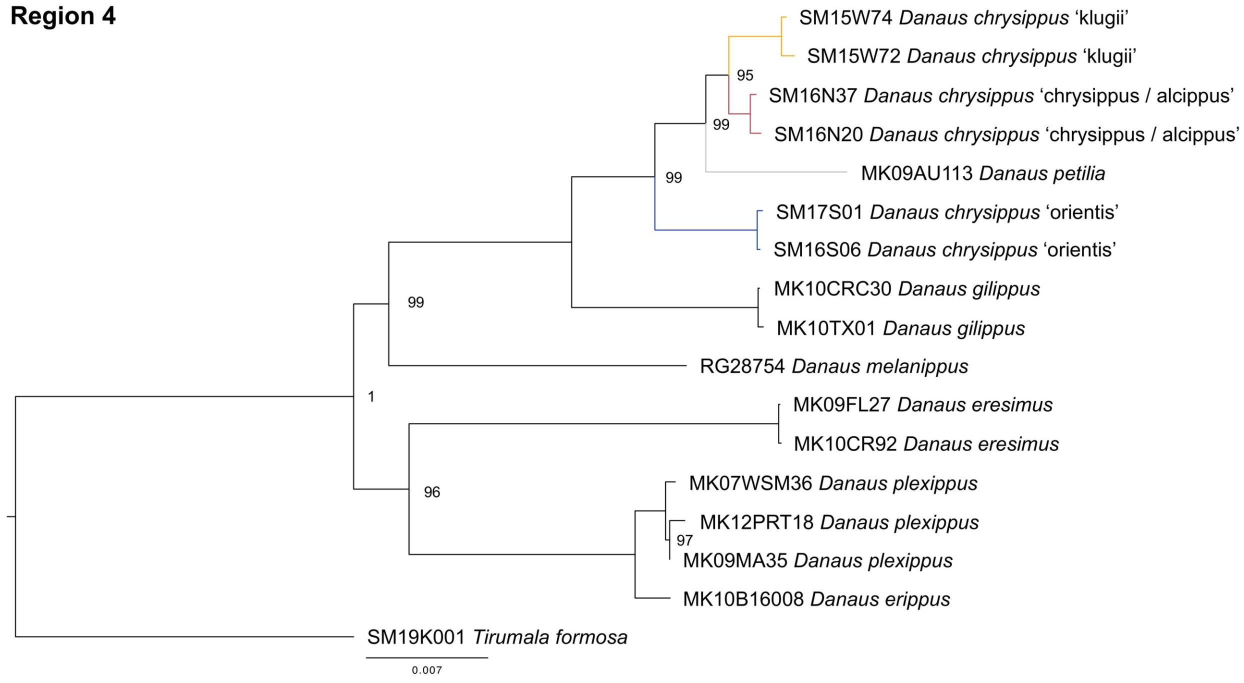


**Figure S8D.** Maximum likelihood phylogenies for concatenated coding sequences for Region 4 (comprising 15 genes). All nodes had bootstrap support of 100% unless otherwise indicated, and nodes with <70% support are shown as polytomies.

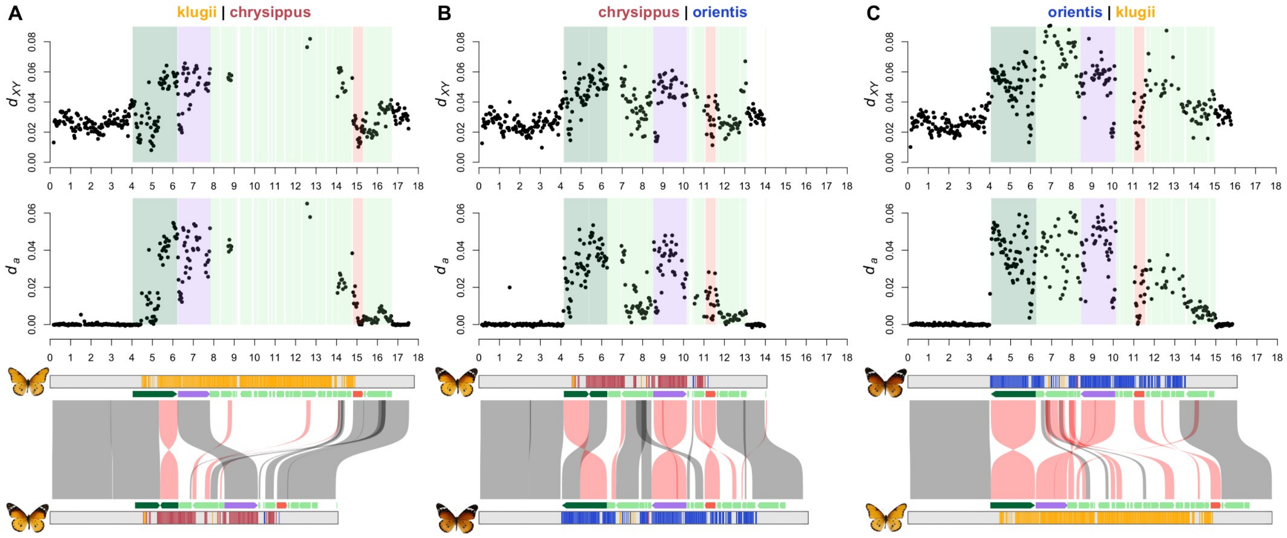


**Figure S9.** Sequence divergence reveals the landscape of recombination suppression in the BC supergene. For each pair of *D. chrysippus* morphs, sequence divergence was computed using reads aligned to one of the two assemblies (the first named morph in each pair). Points in the upper panels represent absolute (*dXY* ) and adjusted (*da*) sequence divergence for 50 kb windows. Tracts corresponding to the four chromosomal Regions (see Figure 1) are shaded. The x-axis is scaled in megabases. The bottom panels show syntenic tracts between pairs of alleles based on pairwise alignments (see Methods for details) syntenic tracts in reverse orientation are shown in red. More stringent alignment parameters were used for these within-species comparisons than the between species comparisons to reduce misalignment between divergent paralogous fragments of Region 3.

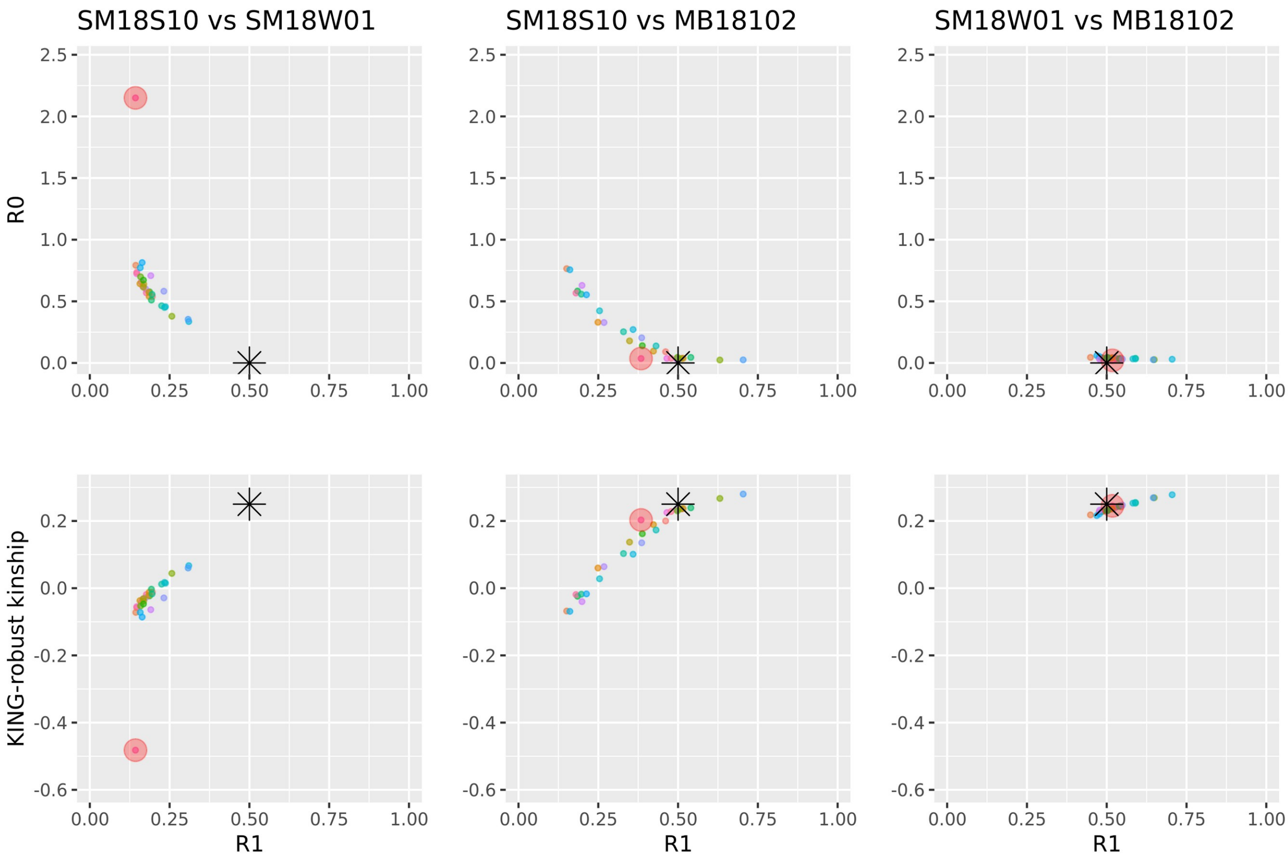


**Figure S10.** The relatedness of autosomal chromosomes between individuals in a trio-binning trial for the brood MB181 reveals that SM18S10 is not the mother of MB18102 and is most likely an aunt. Left column shows aunt (SM18S10) vs father (SM18W01) (unrelated genomes), middle column shows aunt vs F1 (MB18102) (in which some chromosomes show parent-offspring level relatedness and others do not), and right column shows father vs F1 (all chromosomes show parent-offspring level relatedness). The relatedness was calculated based on reads mapped to the *D. plexippus* Dplex_v4 assembly. Small dots represent chromosomes. Red circles show chr15 (labelled as chr7 in the Dplex_v4 assembly). Black stars show the expected position for a parent-offspring relationship. The three measures of relatedness are described by Waples *et al.*[31]. Given that female butterflies do not undergo crossing over, we expected to see two groups of chromosomes in the middle panels: (1) those at which the aunt carries the identical maternal haplotype, which should show parent-offspring relationships; and (2) those at which the aunt carries a different haplotype, which should show values consistent with being unrelated. The observed spread with some chromosomes at intermediate levels suggests that the aunt and mother, which come from a captive stock population, likely had greater than typical relatedness due to inbreeding.

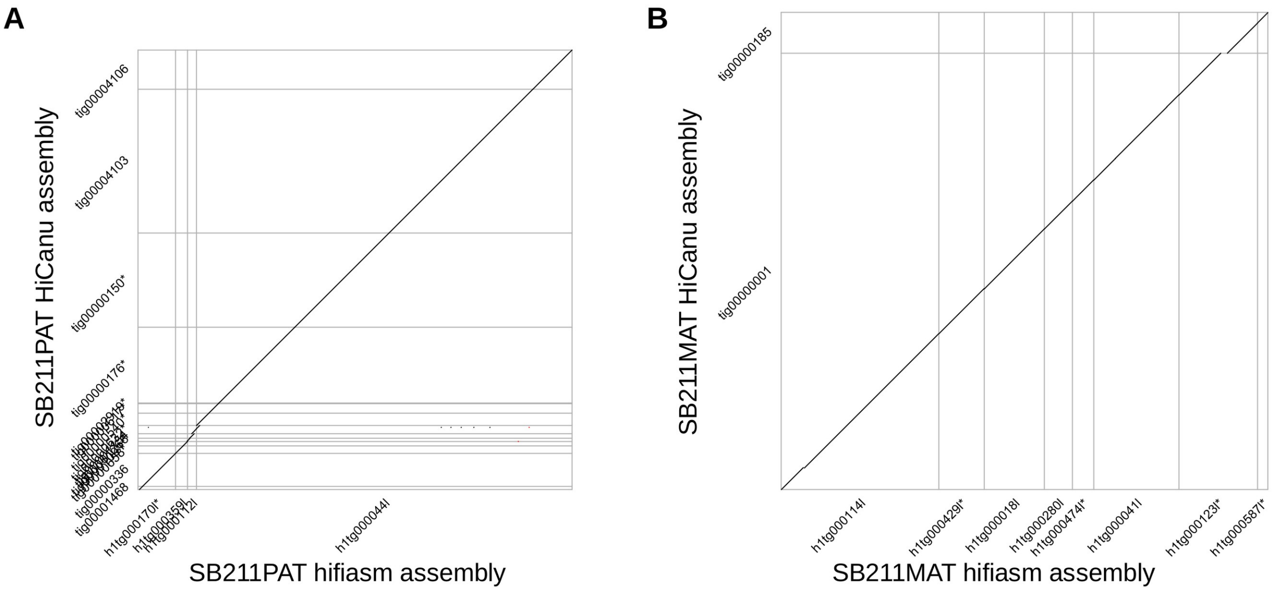


**Figure S11.** Comparison of assemblies made with different assemblers. The paternal (A) and maternal (B) reads from SB211 were each assembled with both hifiasm and HiCanu. To confirm their consistency, minimap2 alignments for all chr15 contigs were visualised. The plots confirm overall consistency between the two assemblers for both assemblies, but also highlight differences in contiguity. The paternal allele (panel A) is more contiguous in the hifiasm assembly (four contigs). The maternal allele (panel B) is more contiguous in the HiCanu assembly (two contigs).
